# Range-wide differential adaptation and genomic vulnerability in critically endangered Asian rosewoods

**DOI:** 10.1101/2023.01.29.524750

**Authors:** Tin Hang Hung, Thea So, Bansa Thammavong, Voradol Chamchumroon, Ida Theilade, Chhang Phourin, Somsanith Bouamanivong, Ida Hartvig, Hannes Gaisberger, Riina Jalonen, David H. Boshier, John J. MacKay

## Abstract

In the billion-dollar global illegal wildlife trade, rosewoods have been the world’s most trafficked wild product since 2005^1^. *Dalbergia cochinchinensis* and *D. oliveri* are the most sought-after rosewoods in the Greater Mekong Subregion^2^. They are exposed to significant genetic risks and the lack of knowledge on their adaptability limits the effectiveness of conservation efforts. Here we present genome assemblies and range-wide genomic scans of adaptive variation, together with predictions of genomic vulnerability to climate change. Adaptive genomic variation was differentially associated with temperature and precipitation-related variables between the species, although their natural ranges overlap. The findings are consistent with differences in pioneering ability and in drought tolerance^3^. We predict their genomic offsets will increase over time and with increasing carbon emission pathway but at a faster pace in *D. cochinchinensis* than in *D. oliveri*. These results and the distinct gene-environment association in the eastern coastal edge suggest species-specific conservation actions: germplasm representation across the range in *D. cochinchinensis* and focused on vulnerability hotspots in *D. oliveri*. We translated our genomic models into a seed source matching application, *seedeR*, to rapidly inform restoration efforts. Our ecological genomic research uncovering contrasting selection forces acting in sympatric rosewoods is of relevance to conserving tropical trees globally and combating risks from climate change.

**Significant statement:** In the billion-dollar global illegal wildlife trade, rosewoods have been the world’s most trafficked wild product since 2005, with *Dalbergia cochinchinensis* and *D. oliveri* being the most sought-after and endangered species in Southeast Asia. Emerging efforts for their restoration have lacked a suitable evidence base on adaptability and adaptive potential. We integrated range-wide genomic data and climate models to detect the differential adaptation between *D. cochinchinensis* and *D. oliveri* in relevance to temperature- and precipitation-related variables and projected their vulnerability until 2100. We highlighted the stronger local adaptation in the coastal edge of the species ranges suggesting conservation priority. We developed genomic resources including chromosome-level genome assemblies and a web-based application seedeR for genomic model-enabled assisted migration and restoration.

## Main

Rosewoods have been the world’s most trafficked wild product since 2005, amounting to 30–40% of the global illegal wildlife trade^1^, which is estimated at 7–23 billion USD annually^4^. *Dalbergia cochinchinensis* Pierre and *D. oliveri* Gamble ex Prain are among the most sought-after and threatened rosewood species. Exploited for their extremely valuable timber^2^, alongside many other valued and threatened tree species in Asia’s tropical and subtropical forests^5^, the growing demand and limited supply have driven prices as high as 50,000 USD per cubic metre^6^. Both these *Dalbergia* species were classified as Vulnerable and Endangered in the 1998 IUCN Red List ^7,8^. The Convention on International Trade in Endangered Species of Wild Fauna and Flora (CITES) has listed the entire *Dalbergia* genus in its Appendix II since 2017 to reduce sequential exploitation of other closely related species^9^. In the IUCN’s latest re-assessment of their endangered status to Critically Endangered in 2022^10,11^, it is suspected that the populations of both species have already experienced a decline of at least 80% over the last three generations, and the decline is likely to continue^12^.

*D. cochinchinensis* and *D. oliveri* are sympatric species, endemic to the Greater Mekong Subregion (GMS) in Southeast Asia, an area of high ecological and conservation concern as 84% of the GMS overlaps with the Indo-Burmese mega biodiversity hotspot^13^. The complex biogeographical and geological histories of the GMS have contributed to its high species richness, heterogeneous landscapes, and high endemism levels^14^. Ancient changes in the distribution of terrestrial and water bodies have been associated with changes in vegetation types and cover^15^. These forests contribute substantially to local livelihoods, economies, food security, and human health^16,17^, though overexploitation undermines their potentially central role to nature-based solutions and most of them are unprotected^4^.

Species- and environment- specific conservation approaches represent an immediate need in response to declining populations^5^. Conservation, collection, and use of genetically diverse germplasm are key to conserving diversity and restoring these rosewood populations. Genetic conservation actions were started in the early 2000s but were limited in scale, usually including fewer than 50 seed-producing trees per country^18–20^. Newer capacity-building initiatives targeting tree nurseries and seed value chain development^21^ may still carry genetic risks associated with the supply and use of germplasm, and may compound the effects of over-exploitation. First, underrepresented genetic diversity during the sourcing of genetic materials can create a genetic bottleneck for the species and reduce the species’ ability to adapt and evolve in a changing climate^22^. Second, mismatch of habitat suitability can result in maladaptation, if populations have strong local adaptation^23^. Third, climate change will likely impose new forces of selection on the current genetic diversity, thus reducing the species’ adaptability, affecting population functioning^24,25^, and leading to increased risk of local extirpations and species’ range collapse^26^. If unaddressed, these risks will reduce both short and long-term effectiveness of restoration projects. The genetic risks call for an understanding of adaptation and its genetic basis in *Dalbergia* species in the GMS to safeguard on-going conservation and restoration efforts. *Dalbergia* are high value species that could be used sustainably and generate income for farmers in developing countries if well-adapted planting material is available^5^. Planting for economic purposes and reducing risks to remaining natural populations of these species seem necessary, where ecological restoration alone is insufficient.

Of the 14,191 vascular plants that are listed as either Vulnerable, Endangered, and Critically Endangered in the IUCN Red List, only 0.1% have their genomes published, far fewer than the 1% reported for listed animals^27^. There is a critical lack of genomic resources in threatened species and a disproportionate representation across taxa, in contrast with the rapid growth in genomic technologies. New reference genomes in threatened species will enable the analysis, of functional genes, higher-resolution studies of species delineation, association mapping and adaptation, genetic rescue, and genome editing^28^. These in turn will help to address important conservation (and restoration) questions such as genetic monitoring of introduced and relocated populations, predicting population viability, disease resistance, synthetic alternatives, and de-extinction^29,30^.

This paper develops an unprecedented understanding of adaptation in critically endangered rosewoods, which integrates genomic analyses, the creation of a novel evidence, and a resource base to inform and expand ongoing conservation efforts. (1) We present genome assemblies of *D. cochinchinensis* and *D. oliveri* at chromosomal and near-chromosomal scale respectively. (2) We analyse range-wide patterns of adaptation by genotyping ~800 trees, and identify differential drivers of adaptive genetic diversity between the two species by using gene-by-environment association analyses. (3) We project current genotypes onto future climate scenarios and predict the potential maladaptation of populations. (4) We deploy an interactive application to predict optimal seed sources, based on our landscape genomic results, in *D. cochinchinensis* and *D. oliveri* for use in restoration under future climate scenarios. Our ecological genomic study in the GMS fills crucial knowledge gaps for genomic adaptation in tropical tree species which are highly underrepresented in the current research literature.

### Chromosome-scale genome characterisation

The *D. cochinchinensis* reference genome assembly (Dacoc_1.4) was 621 Mbp in size comprised of 10 pseudochromosomes (Figure 1a, Supplementary Figure 1, Supplementary Table 1). Whole-genome sequencing of a single seedling of *D. cochinchinensis* produced 165 Gbp (~260 X) long-read data. A diploid-aware draft assembly of 1.3 Gbp with 6,443 contigs and a N50 of 1.35 Mbp was first obtained, with the longest contig between 33.2 Mb at chromosome-arm length. We purged the haplotig and scaffolded the draft genome with 54.97 Gbp (~88.52X) Hi-C chromosome conformation capture reads into 511 scaffolds with a N50 of 60.0 Mb (Supplementary Table 2). The 10 longest scaffolds were considered pseudochromosomes and 98.3% of the contigs were mapped onto them (Figure 1b).

**Figure 1.**
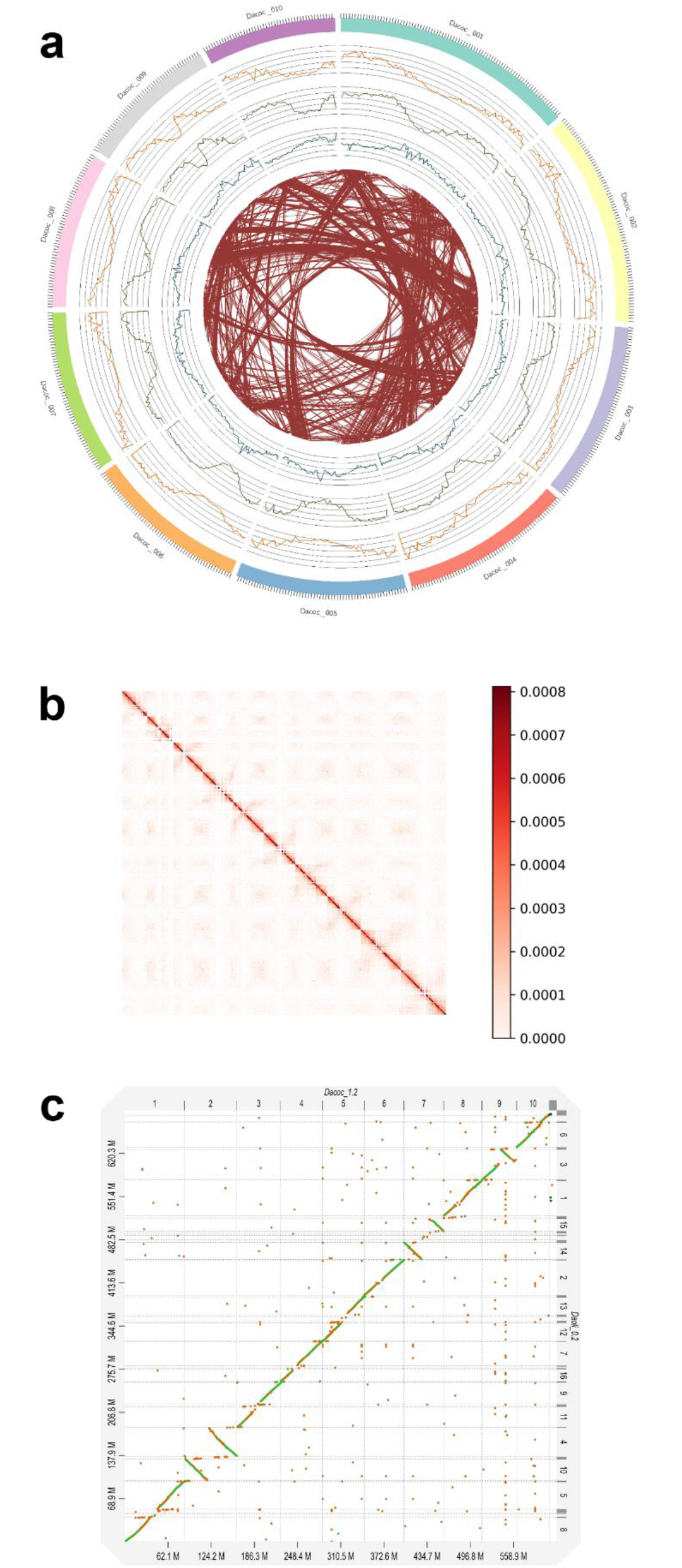
**(a)** Genomic landscape of the 10 assembled pseudochromosomes of D. cochinchinensis (Dacoc_1.4), showing tick marks every 1 Mb, gene density (orange), repeat density (green), 5-mC density (blue), and interchromosomal syntenic arrangement (brown). The densities are calculated in 1-Mb sliding window. **(b)** High-resolution contact probability map of the final D. cochinchinensis genome assembly after scaffolding, revealing the 10 pseudochromosomes at 100 Kbp resolution. **(c)** Syntenic dot plot of assemblies of D. oliveri (Daoli_0.3) against D. cochinchinensis with a minimum identity of 0.25.

The *D. oliveri* draft genome assembly (Daoli_0.3) was 689.25 Mbp in size (Supplementary Figure 1, Supplementary Table 3). Whole-genome sequencing of a single seedling of *D. oliveri* produced 15.13 Gbp (~22X) long-read data. We first obtained a diploid-aware draft assembly of 814.69 Mbp with 3,249 contigs and a N50 of 474.02 Kbp. We purged the haplotig and scaffolded the draft genome with 13.46 Gbp (~20X) Pore-C multi-contact chromosome confirmation capture reads into 2,977 scaffolds with a N50 of 38.43 Mbp. Syntenic analysis of the *D. oliveri* assembly (Daoli_0.3) against the 10 pseudochromosomes obtained in *D. cochinchinensis* (Dacoc_1.4) showed that the 16 largest scaffolds in Daoli_0.3 had 1-to-1 or 2-to-1 correspondences to Dacoc_1.4, implying that Daoli_0.3 was at chromosome-arm length (Figure 1c).

We constructed *de novo* repeat libraries of Dacoc_1.4 and Daoli_0.3, which contained 402 Mbp and 453 Mbp of repeat elements respectively (64.80% and 65.71% of the genomes) (Supplementary Table 4, Supplementary Table 5), the majority of which were annotated as containing LTR elements (46.63% and 48.55%) such as Ty1/Copia (15.25% and 15.75%) and Gypsy/DIRS1 (30.51% and 31.96%). The repeat content of the two genomes was significantly higher than the average among Fabids (~49%), which may be due to the near double amount of LTRs (~22%)^31^.

We predicted and annotated 27,852 and 33,558 gene models in Dacoc_1.4 and Daoli_0.3 respectively, using previous RNA sequencing data (Supplementary Table 6) and protein homology of *Arabidopsis thaliana* and *Arachis ipaensis*. The gene models had a mean length of 4,284.20 and 3942.71 bp respectively, of which 98.3% and 95.5% had an AED score less than 0.5, considered as strong confidence (Supplementary Figure 2). The gene models had a BUSCO v5.1.2 completeness of 96.2% and 88.3% using the eudicots_odb10 reference dataset, with 92.1% and 86.7% being both complete and single copy.

### Range-wide genomic scan for adaptive signals

We obtained initial pools of 1,832,629 and 3,377,855 SNPs from genotyping 435 and 331 individuals of *D. cochinchinensis* and *D. oliveri* respectively, across their natural ranges (Supplementary Table 7), and final pools of 180,944 and 193,724 SNPs after filtering for missing data, minimum allele frequency, and linkage disequilibrium. The samples represented previous sampling work^32,33^ and new sampling that covered all known existing populations.

We employed the sparse non-negative matrix factorisation (sNMF) algorithm to determine the optimal number of ancestral populations (K) for *D. cochinchinensis* and *D. oliveri* as 13 and 14 respectively (Supplementary Figure 3, Supplementary Figure 4, Supplementary Figure 5). These results were much higher than the previous estimation of K = 5 – 9 for the same species using nine microsatellite markers and 19 SNPs^32,33^. The analysis revealed a highly resolved hierarchical genetic structure for both species and distinct population clusters around the Cardamon Mountains in southwest Cambodia and in northern Laos. Our calculation gave a larger genomic inflation factor (λ) in *D. cochinchinensis* (range from 0.071 (evapotranspiration) to 0.25 (precipitation of driest quarter), mean of 0.13, standard deviation of 0.049) than that in *D. oliveri* (range from 0.038 (evapotranspiration) to 0.081 (mean diurnal range), mean of 0.056, standard deviation of 0.016 (Supplementary Table 8).

The numbers of SNPs found to be adaptive for at least one of the environmental variables were 20,373 (11.3%) and 6,953 (3.59%) in *D. cochinchinensis* and *D. oliveri* respectively (| *Z*-value | > 2 & *Q*-value < 0.01), after correcting for population structure (optimal K) and genomic inflation (Supplementary Figure 6, Supplementary Figure 7, Supplementary Table 9). Relatively few SNPs were associated with all or many environmental variables; 4 SNPs were associated with 11 out of 13 variables tested in *D. cochinchinensis*, and 46 SNPs were associated with all 12 variables in *D. oliveri*. These findings revealed the complex and polygenic nature of environmental adaptation, where multiple forces of natural selection can act together via different environmental cues and affect overlapping loci.

In *D. cochinchinensis*, ‘precipitation in the driest quarter’ was the environmental variable (wc2.1_30s_bio_17) and the strongest gene-environmental association with a SNP on chromosome 3 at position 36,345,659 (LFMM Z = 6.07237, *Q* = 4.77e-29). The SNP was located within the gene Dacoc08834, a homologue of the Ubiquitin-like-specific protease 1B ULP1B. The highest allele frequencies of this SNP were found in the southwest of Cambodia with the highest precipitation of the driest quarter (Figure 2). ULP1B is one of the ubiquitin like-specific proteases that mediate the maturation and deconjugation of a small ubiquitin-like modifier (SUMO) from target proteins as part of post-translational modification^34^. The SUMO process in plants has been shown to regulate stress responses including to drought, heat, salinity, and pathogens^35–37^ and timing of flower initiation^38^, which might explain the strong association with the drought stress associated with the said environmental factor. In an analysis of transcriptomes from 6 *Dalbergia* species, ubiquitin-related proteins were found to be overrepresented compared to other legumes^27^. Taken together, these observations suggest that ubiquitin-related proteins have a role in *Dalbergia* adaptation to water assimilation.

**Figure 2.**
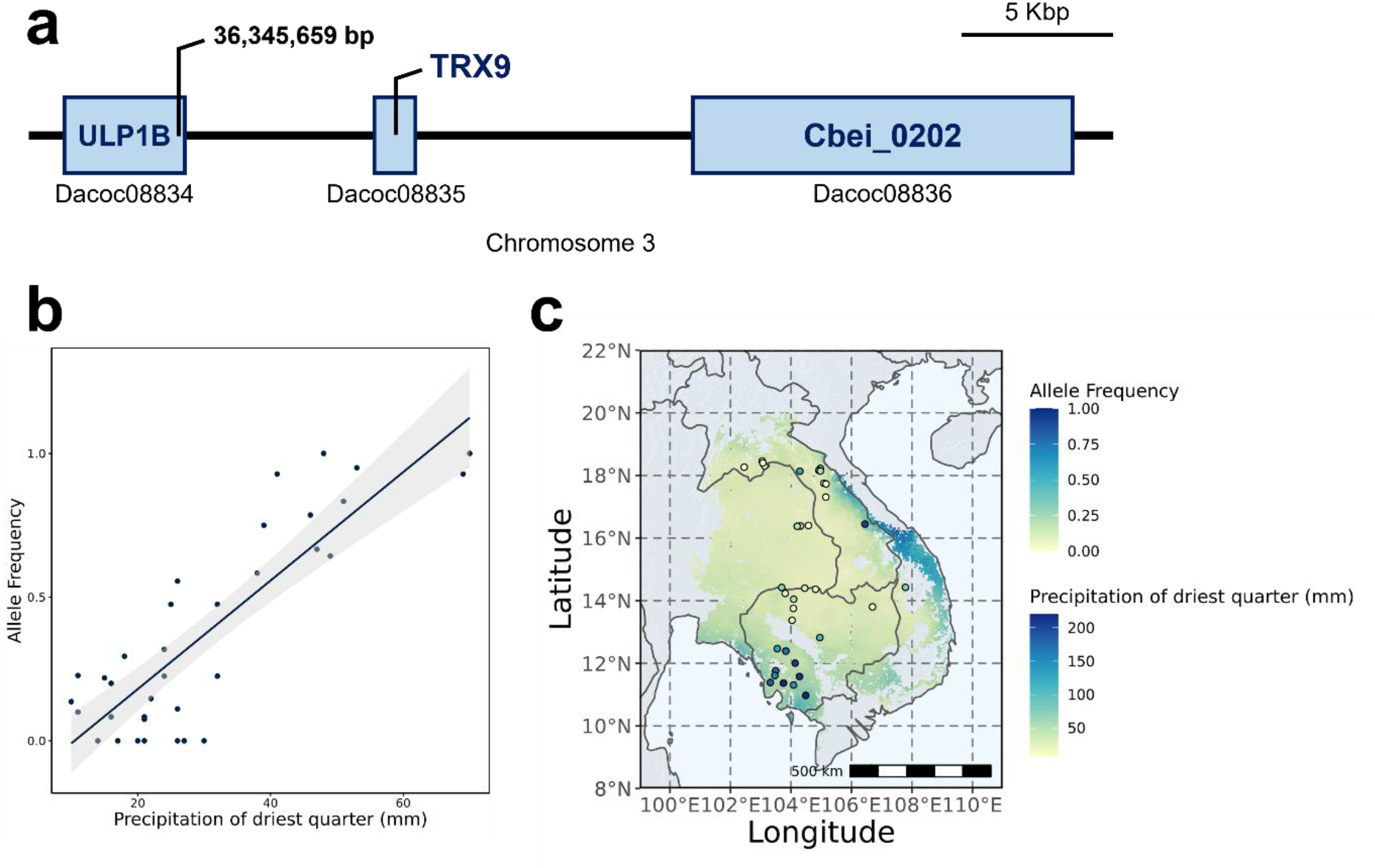
**(a)** The most significant gene-environment association at 36,346,659 bp on chromosome 3, within the Dacoc08834 gene and upstream of Dacoc08835 and Dacoc08836 genes, which are homologues of ULP1B, TRX9, and Cbei_0202 respectively. **(b)** and **(c)** Correlation between allele frequency and wc2.1_30s_bio_17 (Precipitation of driest quarter) for this locus.

By contrast, the strongest association in *D. oliveri* was between precipitation of the wettest quarter (wc2.1_30s_bio_16), and a SNP on the scaffold Daoli_0035 at the position 107,725 (LFMM Z = 6.1895, *Q* = 6.36e-102). The locus was 3,254 bp upstream of a predicted gene model Daoli32516 and 5,010 bp downstream of the gene Daoli32517, a homologue of tatC-like protein YMF16.

### Differential adaptation related to temperature and precipitation

Isothermality (wc2.1_30s_bio_3) was identified as the most important overall driver of both neutral and adaptive genomic variation among non-spatial environmental variables in *D. cochinchinensis*, based on our gradient forest (GF) model (Figure 3, Supplementary Figure 8a), in contrast to ‘precipitation of the wettest quarter’ (wc2.1_30s_bio_16) in *D. oliveri* (Figure 4, Supplementary Figure 8b). Spatial variables, as principal coordinates of a neighbourhood matrix (PCNM), were the most important variables that explained both neutral and adaptive genomic variation, which was unsurprising given strong isolation by distance was known in these species^32^ and environmental adaptation only affects a small portion of the genome^39^. Soil factors were among the lowest ranked variables for gene-environment associations for both species. We observed different patterns of geographic variation in *D. cochinchinensis* and *D. oliveri* when fitting the GF models across their native ranges. *D. cochinchinensis* had strong differentiation between North and South populations at around 16°N, that was mainly driven by isothermality (wc2.1_30s_bio_3) as seen in the PCA loadings. On the other hand, *D. oliveri’s* major differentiation was between coastal and inland areas, driven by both precipitation of the wettest quarter (wc2.1_30s_bio_16) and mean diurnal range (wc2.1_30s_bio_2). The eastern coastal areas in Vietnam showed particularly strong differences in environmental associations with adaptive variation and neutral variation for both *D. cochinchinensis* and *D. oliveri* (Figure 5).

**Figure 3.**
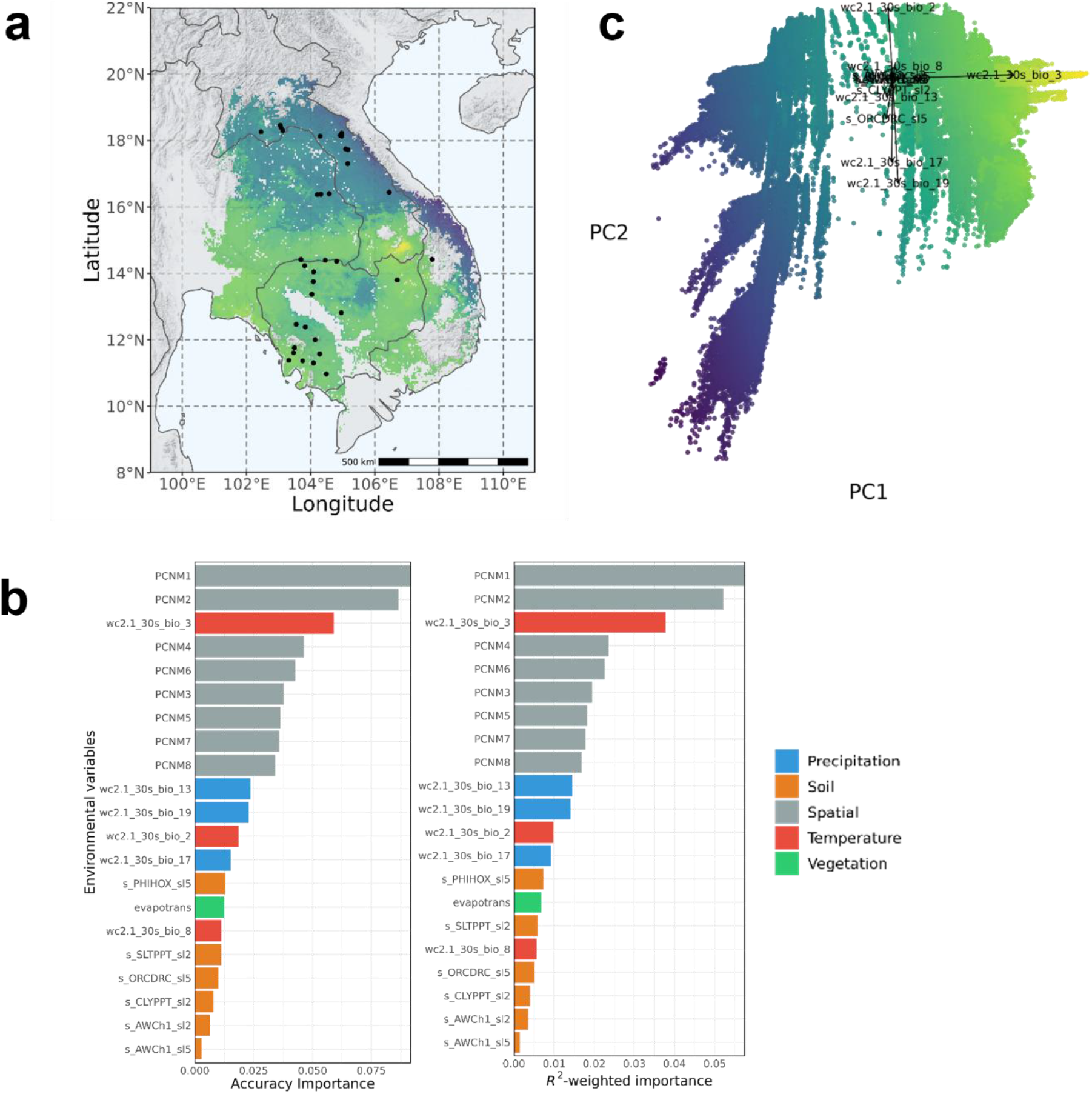
**(a)** Adaptive genomic variation across the species range predicted by GF model for D. cochinchinensis, visualised using the first two principal axes from the PCA. **(b)** Accuracy and R^2^-weighted importance for environmental predictor variables which explained adaptive genomic variation (adaptive SNPs) by the GF model. **(c)** Principal component analysis (PCA) of the adaptive genomic variation predicted by the GF model across the species range. Loadings are the environmental factors.

**Figure 4.**
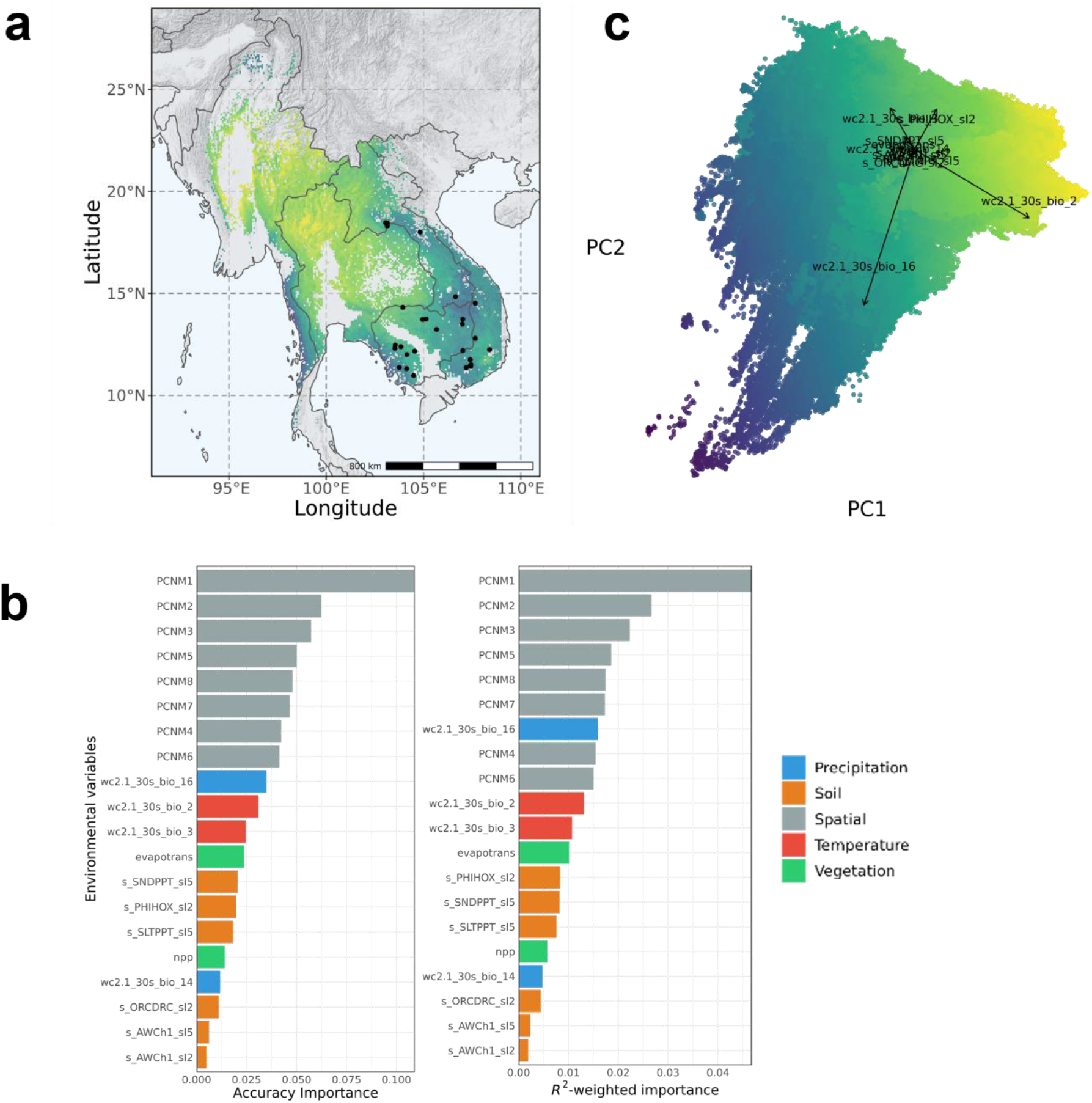
**(a)** Adaptive genomic variation across the species range predicted by GF model for D. oliveri, visualised using the first two principal axes from the PCA. **(b)** Accuracy and R^2^-weighted importance for the environmental predictor variables which explained the adaptive genomic variation (adaptive SNPs) by the GF model. **(c)** Principal component analysis (PCA) of the adaptive genomic variation predicted by the GF model across the species range. The loadings are the environmental factors.

**Figure 5.**
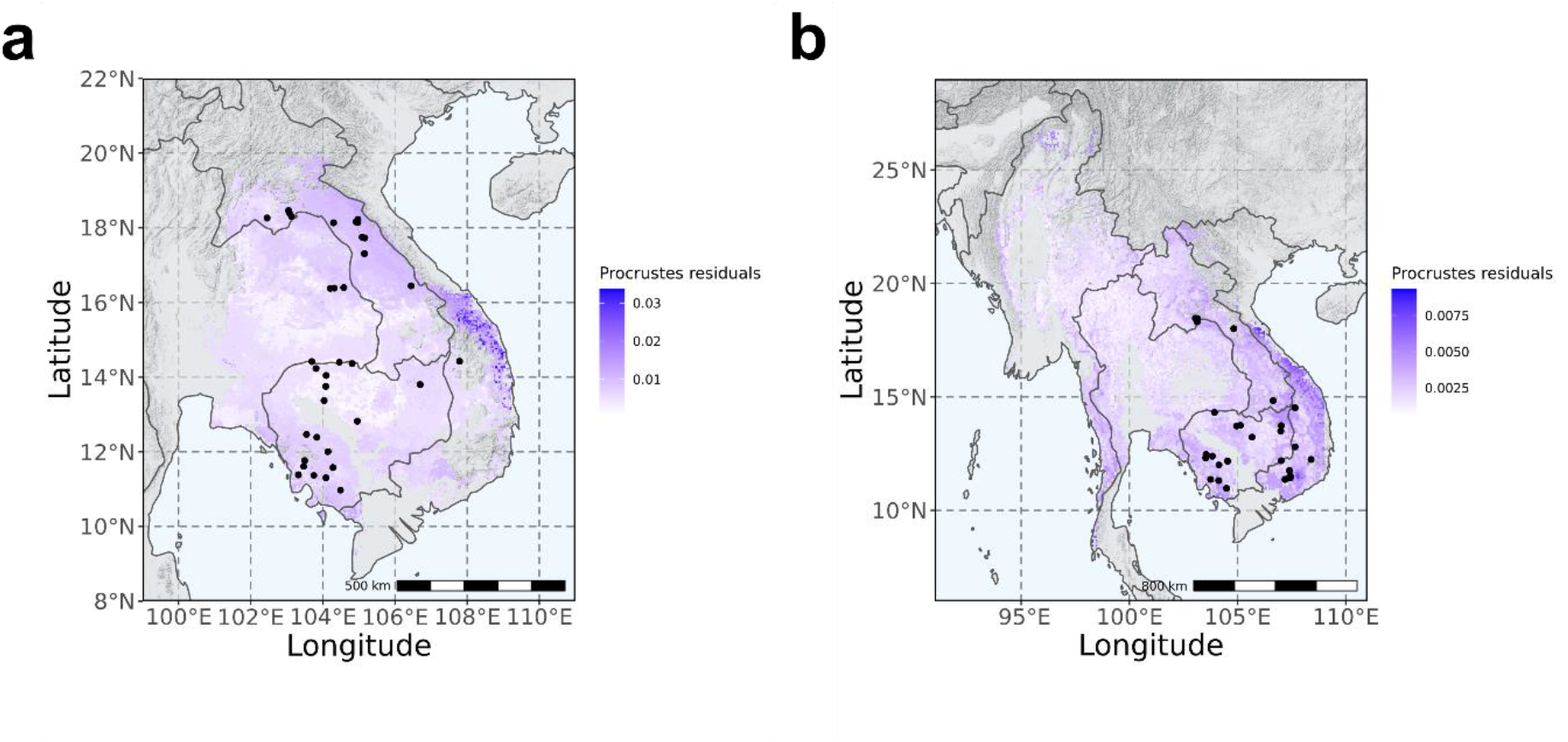
Procrustes residuals between neutral and adaptive gene-environmental associations for (a) D. cochinchinensis and (b) D. oliveri.

We compared the allelic frequency turnover functions of the neutral and adaptive genomic variation for each environmental predictor variable. Adaptive genomic variation was significantly more strongly associated with environmental gradients than neutral variation (Supplementary Figure 9). There was only one exception, where available soil water capacity at a depth of 60 cm (s_AWCh1_sl5) was near-zero but of similar importance in explaining neutral and adaptive variation, regardless of the environmental gradient.

When exposed to drought stress under controlled conditions, *D. cochinchinensis* was more anisohydric than *D. oliveri*, which means that *D. cochinchinensis*, as a pioneering species with faster growth, optimises carbon assimilation and better tolerates reduced water availability^3^. *D. oliveri* is often found in moist areas and along streams and rivers^40^, and the morphological characteristics of its seeds suggest that secondary dispersal by water is likely^32^. This could explain how isothermality, which is a useful metric in tropical environments^41^ and shown to influence plant height growth^42^, had a dominant effect in the adaptive variation only in *D. cochinchinensis*. Pioneering species maximise height growth in early successional habitats to meet their light requirements^43^, consistent with the observation of higher photosynthetic pigment levels in *D. cochinchinensis*^3^. On the other hand, the effect of precipitation of the wettest quarter could act on selection in seed dispersal and survival in *D. oliveri* in the wet season. Temperature and precipitation, and their variability such as isothermality^44^ have been widely reported as the most important drivers shaping patterns of productivity and adaptation in tree species across the world^45–47^.

To fill the current gaps in existing conservation actions, populations that are underrepresented but display distinct adaptive variation should be prioritised to avoid the potential loss of unique genetic diversity. Populations at the edge of the species ranges should be prioritised based on our findings on adaptive variation showing their distinct allelic frequencies and adaptation; however, they are currently underrepresented in conservation efforts and existing protected area networks. Importantly, hotspots of differential adaptive variation near the edges of species ranges are shared between *D. cochinchinensis* and *D. oliveri*. This observation reinforces the role of marginal populations in preserving evolutionary potential for range expansion and persistence due to their adaptation to distinct environmental conditions^48^.

### Genomic vulnerability under different climate change scenarios

Genetic offset in the form of Euclidean distance represented the mismatch between current and future gene-environment association, which was modelled over five general circulation models (GCMs), namely MIROC6, BCC-CSM2-MR, IPSL-CM6A-LR, CNRM-ESM2-1, MRI-ESM2-0, under WCRP CMIP6 (Supplementary Figure 10). For both *Dalbergia* species, genetic offset generally increased over time (*P* = 2.71e–10) and shared socioeconomic pathway (*P* = 4.54e–14), which implies increased carbon emission (Figure 6a, Supplementary Table 10). However, *D. cochinchinensis* shows a significantly larger increase in genetic offset over time compared to *D. oliveri* (*P* = 0.025), suggesting that *D. cochinchinensis* is more susceptible to any mismatch of current genotypes and future climate. The geographic patterns of genetic offset also differed between the two species: *D. cochinchinensis* had an increasing offset across the entire range, while *D. oliveri* had a distinctly high offset in the southeast part of the range (Figure 6b–c). The variation in genomic offset between two species was mainly driven by the strong association with isothermality (wc2.1_30s_bio_3) in *D. cochinchinensis*, as demonstrated in the GF model, as it contributed to ~75% of the genomic offset on average (Figure 6d). Isothermality had a smaller effect (~35%) in *D. oliveri* (Figure 6e).

**Figure 6.**
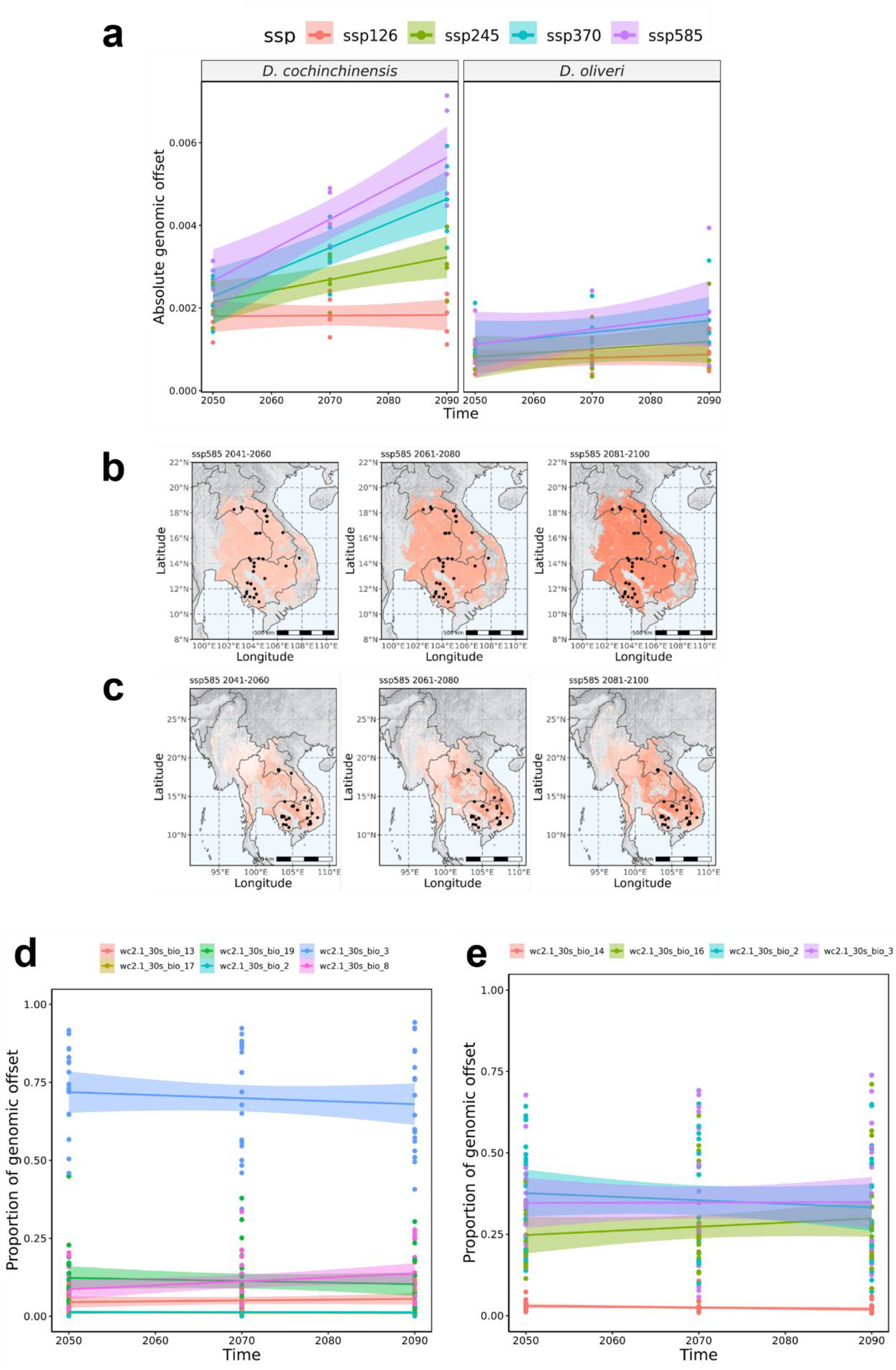
**(a)** Absolute genomic offset of gene-environment association, quantified as the Euclidean distance, of D. cochinchinensis and D. oliveri in 4 SSPs (126, 245, 370, and 585) over three bidecades (2041–2060, 2061–2080, 2081– 2100) averaged across five GCMs (BCC-CSM2-MR, CNRM-ESM2-1, IPSL-CM6A-LR, MIROC6, MRI-ESM2-0). Scaled genomic offset across the range of **(b)** D. cochinchinensis and **(c)** D. oliveri, using SSP585 between 2041 and 2060 as an example. Proportion of genomic variation explained by environmental variables in **(d)** D. cochinchinensis and **(e)** D. oliveri.

Our prediction contrasts with a separate sensitivity-and-exposure modelling study which predicted that *D. oliveri* is likely to be slightly more vulnerable to climate change by 2055 (2041–2070 period) than *D. cochinchinensis*^12^. It used growth rate and seed weight as proxy traits, predicting that both species have equally high sensitivity to climate change, but that *D. oliveri* is more exposed to the threat. Our findings predict that the dominant environment factor of isothermality could give more weight to the species’ vulnerability. As discussed, isothermality is likely to affect the productivity and growth in pioneering species like *D. cochinchinensis* more than later successional species like *D. oliveri*. Our work supports that isothermality and other temperature variation factors will serve as more reliable indicators to predict the climate response of *D. cochinchinensis* and encourages further studies of this response, such as greenhouse or common garden experiments to validate the prediction with empirical data.

The different geographical patterns of genomic vulnerability support species-specific recommendations in conservation and restoration. While climate change is likely to affect *D. cochinchinensis* evenly across its range, greater attention is needed on the representation of adaptive variation in germplasm collection and conservation units; sampling should target edge populations in particular as they show potential signals of local adaptation, where the environmental associations between adaptive and neutral variation are the greatest. By contrast, we recommend targeting hotspots of vulnerability in *D. oliveri*, especially around the borders between Cambodia, Laos, Vietnam, and Thailand, to improve conservation efforts.

In a rapidly changing environment, forest trees either persist through migration or phenotypic plasticity, or will extirpate^45^ when environmental change outpaces adaptation potential. The spatially explicit model of genomic vulnerability helps to develop conservation decisions balancing between *in situ* adaptation and assisted migration, as populations with lower vulnerability are likely to persist through adaptation^49^.

### Genomic model-enabled assisted migration and restoration

We developed *seedeR*, an open-source web application that is freely available from https://trainingidn.shinyapps.io/seedeR/, where users can input the species (*D. cochinchinensis* or *D. oliveri*), shared socioeconomic pathways (SSP), time period, and geographical coordinates of the target restoration or planting site. With these inputs, *seedeR* predicts the genomic similarity between a current germplasm source and target site from allelic frequency turnover functions and genetic offset and projects them onto the species range. We demonstrate the utility of *seedeR* for a hypothetical target restoration site (106° N, 14° E) in northeast Cambodia for both *D. cochinchinensis* and *D. oliveri*, under the future climate scenario of SSP370 between 2081 and 2100 (Figure 7). In both predictions, the genomic similarity was the highest at proximity to several hundreds of kilometres and decreased when further away. Commonly, coastal regions in northeast Vietnam, which were predicted to have the strongest local adaptation in both species, showed a lower genomic similarity. The geographical scale of suitable seed sources has an important implication as too many forest landscape projects collect seeds from very close (a few kilometres) to restoration sites to feed the “local is best” paradigm^50^, while our predictions showed otherwise. It is also important to note that local tree populations in landscapes in need of restoration are often degraded and have low genetic diversity. Genetic quality of seed should be ensured by collecting seed from large populations and many unrelated trees, even if this means collecting from trees at distances much further from the target restoration site.

**Figure 7.**
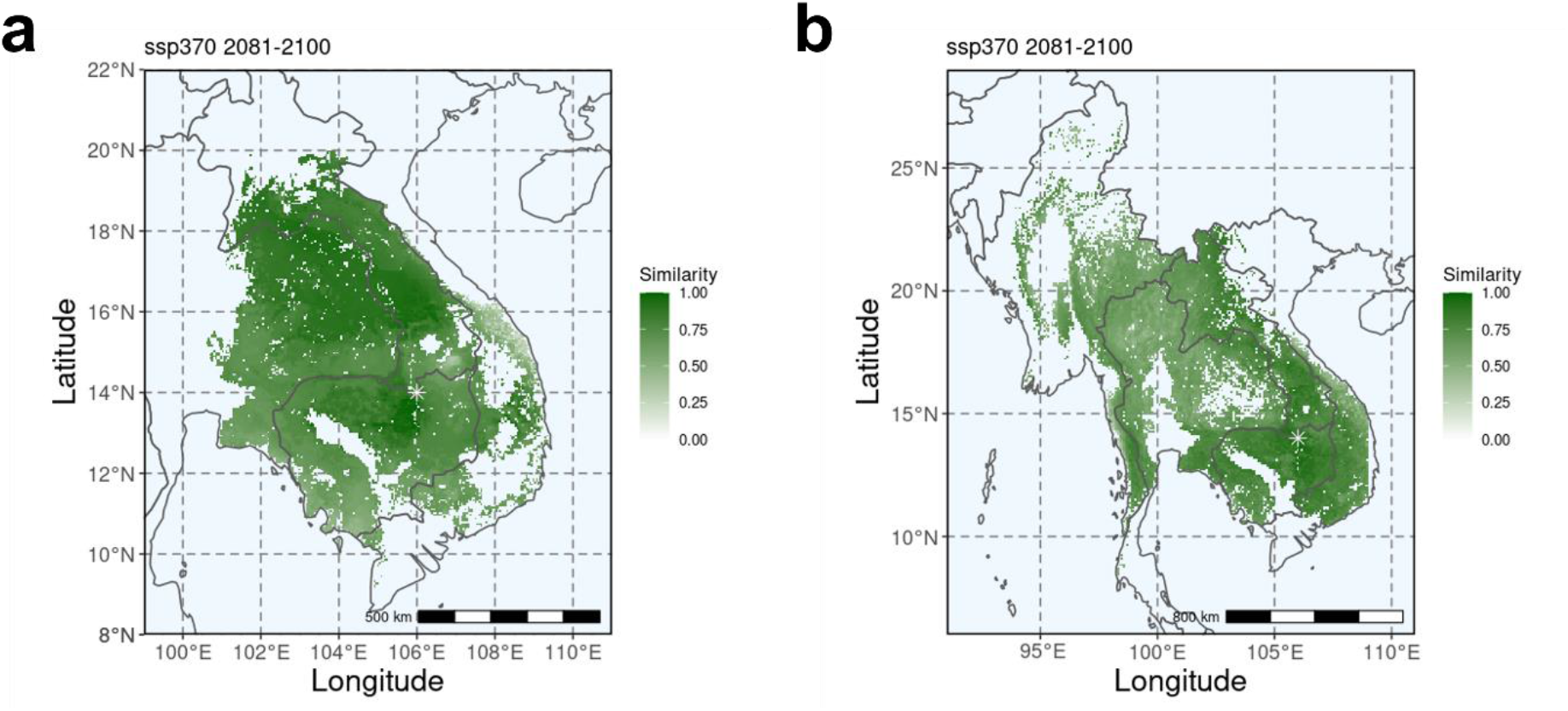
Genomic similarity (scaled between 0, most dissimilar, and 1, most similar) between a hypothetical future restoration site (106° N, 14° E) and the current potential germplasm sources under the future climate scenario of SSP370 between 2081 and 2100 for **(a)** D. cochinchinensis and **(b)** D. oliveri predicted on seedeR (https://trainingidn.shinyapps.io/seedeR/).

Matching seed sources and restoration sites remains one of the keys for effective conservation and restoration^51^, in line with the importance of adaptive variation and potential in genetic materials. Our genome-enabled prediction tool considers the future climate of restoration sites, which in turn will greatly influence the future resilience and productivity of these species. In the case of maladaptation and extirpation due to environmental change^52^, when the classical preference for local provenance may no longer hold, deliberate transfer of germplasm along climate gradients may be necessary^53^. Especially in the case of *Dalbergia*, when many local populations have extirpated or are very small in size, and large environmental association was predicted, assisted migration based on admixture and predictive provenancing are deemed more appropriate for the species to facilitate adaptation of the populations under climate change^54^. Genetic materials from regions with strong adaptive genomic variation, such as coastal Vietnam, can be moved to suitable regions using the *seedeR* prediction to facilitate gene flow and maintain unique genetic components of the population by admixture^53^. Hotspots of vulnerable populations such as those in northern Cambodia are suitable to be moved to new suitable areas to prevent loss of genetic diversity.

The *seedeR* application helps to visualize these spatially explicit predictive models of genomic vulnerability and match, which are most useful to frontline practitioners and managers^55^. Not only can it inform conservation and management strategies, but by simplifying the analytical pipelines through a user-friendly platform, it will also directly reduce the gap between conservation and genomics; a challenge faced for dissemination of genomic knowledge^56^.

### Narrowing the gap between conservation and genomics

Our study characterises range-wide gene-environment association in two sympatric endangered species, *D. cochinchinensis* and *D. oliveri*, for which there was virtually no prior knowledge on adaptability. Building on previous understanding of their different physiologies, we demonstrate their differential adaptive characteristics, which point to species-specific implications for their conservation. These findings on differential genomic adaptation between sympatric species sheds novel understanding on tropical forests, which in particular harbour many threatened species, at risk from threats associated with climate change.

We show how genomic technologies can directly support rapid decision-making and conservation activities. The separation between scientific and conservation communities represents a long-standing challenge, such that advances in scientific research and specifically genomic technologies are often inaccessible to the conservation side, which hinders translational science^57,58^. Through engagement with diverse stakeholders and conservation activities, we were strongly motivated to deliver the results of this study in a user-friendly (e.g. *seede*R) and spatially explicit manner that can be integrated with ongoing conservation work.

## Methods

### Plant materials and sample preparation for genome assemblies

Dried seeds of *Dalbergia cochinchinensis* and *D. oliveri* were collected from the Bolikhamxay, Khamkend, Laos, and Phnom Penh, Cambodia in 2018 by their forestry authorities respectively. We germinated the seeds in a greenhouse at 30°C with 16L/8D photoperiod. Leaf tissues were harvested from a selected 1-year-old individual for each species and ground in liquid nitrogen with a mortar and pestle.

High-molecular-weight genomic DNA was extracted from the reference individual with Carlson lysis buffer (100 mM Tris-HCl, pH 9.5, 2% CTAB, 1.4 M NaCl, 1% PEG 8000, 20 mM EDTA) followed by purification using the QIAGEN Genomic-tip 500/G. The quantity and quality of genomic DNA were determined with NanoDrop 2000 (Thermo, Wilmington, United States) and Qubit 4 (Thermo Fisher Scientific, United Kingdom). DNA integrity was preliminary assessed with a 0.4% agarose gel against a NEB Quick-Load® 1 kb Extend DNA Ladder. A DNA sample passed the quality check only when a single band could be mapped near a lambda DNA band (~ 48.5 kb).

### Genomic sequencing and assembly of D. cochinchinensis

For Oxford Nanopore sequencing, 9 μg of extracted DNA was size-selected using the Circulomics Short Read Eliminator XL Kit (Maryland, United States) to deplete fragments < 40 Kbp. Three libarires were prepared each starting from 3 μg of size-selected DNA was used in each library preparation with the Oxford Nanopore Technologies Ligation Sequencing Kit (SQK-LSK110). The libraries were sequenced on two R10.3 (FLO-109D) flow cells on a GridION sequencer for ~ 72 hours. Real-time basecalling was performed in MinKNOW release 19.10.1. Raw reads with Phred score lower than 8 were filtered.

For PacBio sequencing, DNA samples were sent to the Genomics & Cell Characterization Core Facility at the University of Oregon for DNA library preparation and sequencing. Throughout the sample preparation, the quality of DNA was assessed using Fragment Analyzer 1.2.0.11 (Agilent, United States). 20 μg of unsheared genomic DNA was used for library preparation using the SMRTbell Express Template Prep Kit 2.0 (Pacific Biosciences, United States). The library was size selected using the BluePippin system (Sage Science, United States) at 45 kb and then sequenced on a single SMRT 8M cell on a Sequel II System (2.0 chemistry) using the Continuous Long-Read Sequencing (CLR) mode with a movie time of 30 hours.

For Hi-C sequencing, we harvested 0.5 g of fresh leaf from the same reference individual and immediately cross-linked the finely chopped tissue in 1% formaldehyde for 20 minutes. The cross-linking was then quenched with glycine (125 mM). The cross-linked samples were ground in liquid nitrogen with a mortar and pestle and shipped to Phase Genomics (Seattle, USA) for library preparation and sequencing. The Hi-C library was prepared with the restriction enzyme DpnII, proximity-ligated, and reverse-crosslinked using Proximo Hi-C Kit (Plant) v2.0 (Phase Genomics, Seattle, USA). The library was sequenced on a HiSeq4000 for ~300 M 150-bp paired-end sequencing.

### Genomic sequencing of D. oliveri

For Nanopore sequencing, the same protocol and procedure were used as for *D. cochinchinensis* (see above).

For Pore-C sequencing, the library was prepared with the protocol and reagents described by Belaghzal et al.^59^ with minor modifications. We harvested 2 g of fresh leaf from the same reference individual as for the Nanopore library and immediately cross-linked the finely chopped tissues in 1% formaldehyde for 20 minutes. The cross-linking was quenched with 125 mM glycine for 20 minutes and then the samples were ground in liquid nitrogen with a mortar and a pestle. Cell nuclei were isolated with a buffer containing 10 mM Trizma, 80 mM KCl, 10 mM EDTA, 1 mM spermidine trihydrochloride, 1 mM spermine tetrahydrochloride, 500 mM sucrose, 1% (w/v) PVP-40, 0.5% (v/v) Triton X-100, and 0.25% (v/v) β-mercaptoethanol, and then passed through a 40 μm cell strainer. The suspension was centrifuged at 3,000 g, according to the estimated genome size of ~ 700 Mbp. Chromatin was denatured with the restriction enzyme NlaIII at a final concentration of 1 U/μL (New England Biolabs, United Kingdom) at 37°C for 18 hours. The enzyme was heat-denatured at 65°C for 20 minutes at 300 rpm rotation in a thermomixer. Proximity ligation, protein degradation, decrosslinking, and DNA extraction were performed according to the original Belaghzal protocol. The Pore-C library was prepared with the Oxford Nanopore Technologies Ligation Sequencing Kit (SQK-LSK110), then sequenced on two R10.3 (FLO-109D) Nanopore flow cells on a GridION sequencer for ~ 72 hours. The flow cell was washed once every 24 hour with the Flow Cell Wash Kit (EXP-WSH003).

### Assembly pipelines

Raw reads shorter than 500 bp were filtered. Due to the heterozygous nature of the wild individual, we assembled the sequences with Canu 2.1.1 using the options “corOutCoverage=200 correctedErrorRate=0.16 batOptions=−dg 3 -db 3 -dr 1 -ca 500 -cp 50”. We then used purge_haplotigs v1.1.1 to collapse the assembly by separating the primary assembly and haplotigs.

Hi-C reads (for *D. cochinchinensis*) were mapped to the draft genome assembly using hicstuff 2.3.2^60^ to generate the contact matrix, which was then used to scaffold and polish the assembly using instaGRAAL 0.1.2^61^ with default options to produce the final assembly Dacoc 1.4 after removing contamination.

Pore-C reads (for *D. oliveri*) were mapped to the draft genome assembly and used to generate contact map with the Pore-C-Snakemake (https://github.com/nanoporetech/Pore-C-Snakemake) and produce a merged_nodups (.mnd) file, which contains a duplicate-free list of paired alignments from the Pore-C reads to the draft assembly. The draft assembly and the merged_nodups file were used for scaffolding in 3D-DNA (version 180419) and produce the final genome Daoli 0.3.

To validate the scaffold arrangement, Daoli 0.3 was aligned to that of *D. cochinchinensis* (Dacoc 1.4) using minimap2 and D-GENIES^62^ to produce a dot plot for visualising similarity, repetitions, breaks, and inversions, with a minimum identity of 0.25.

### De novo repeat library

A *de novo* repeat library was constructed using RepeatModeler 2.0.1^63^, which incorporated RECON 1.08^64^, RepeatScout 1.0.6^65^, and TRF 4.0.9^66^ for identification and classification of repeat families. We then used RepeatMasker 4.1.1^67^ to mask low complex or simple repeats only (“-noint”). A *de novo* library of long terminal repeat (LTR) retrotransposons was constructed on the simple-repeat-masked genome using LTRharvest^68^ and annotated with the GyDB database and profile HMMs using LTRdigest^69^ module in the genometools 1.6.1 pipeline. Predicted LTR elements with no protein domain hits were removed from the library. We applied the RepeatClassifier module in RepeatModeler to format both repeat libraries. We merged the libraries together and clustered the sequences that were ≥ 80% identical by CD-HIT-EST 4.8.1^70^ (“-aS 80 -c 0.8 -g 1 -G 0 -A 80”) to produce the final repeat library.

### Gene models and annotation

Filtered mRNA-sequencing data for *D. cochinchinensis* (50.5 Gbp) and *D. oliveri* (54.4 Gbp) from a previous project^27^ (NCBI Bioproject: PRJNA593817) were aligned against the genome assembly using STAR v2.7.6 and assembled using the genome-guided mode of Trinity v2.13.2. Protein sequences were obtained from *Arabidopsis thaliana* (Araport11)^71^ and *Arachis ipaensis* (Araip1.1)^72^. After soft-masking the genome with the *de novo* repeat library using RepeatMasker (Dfam libraries 3.2), the transcript and protein evidences were used to produce gene models using MAKER 3.01.03^73^. The MAKER pipeline was iteratively run for two more rounds to produce the final gene models. In between each run of MAKER, the gene models were used to train the *ab initio* gene predictors SNAP (version 2006-07-28)^74^ and AUGUSTUS 3.3.3^75^ which were used in the MAKER pipeline. tRNA genes were predicted with tRNAscan-SE 1.3.1^76^. The quality of the gene models was assessed with two metrics: the annotation edit distance (AED) in MAKER 3.01.03^73^ and the BUSCO score (v5.1.2)^77^.

### Population sampling

We obtained a collection of 435 and 331 foliage samples of *Dalbergia cochinchinensis* and *D. oliveri* from 35 and 28 localities across their native range (Supplementary Table 11). These samples were a combination of those collected in a previous study^32^ and newly between 2019 and 2020. Genomic DNA was purified using a two-round modified CTAB protocol (2% CTAB, 1.4 M NaCl, 1% PVP-40, 100 mM Tris-Cl pH 8.0, 20 mM EDTA pH 8.0, 1% 2-mercaptoethanol) with sorbitol pre-wash (0.35 M Sorbitol, 1% PVP-40, 100 mM Tris-Cl pH 8.0, and 5 mM EDTA pH 8.0) as the samples were rich in polyphenols and polysaccharides^78^. Genomic DNA was treated with 5 μL RNase (10 mg/mL). Quality and quantity of the genomic DNA were assessed using NanoDrop One (Thermo, Wilmington, United States) and Qubit dsDNA BR Assay kit on Qubit 4 (Thermo, Wilmington, United States) respectively.

### Genotyping-by-sequencing (GbS)

DNA samples were normalised to 200 ng suspended in 10 μL water and sent to the Genomic Analysis Platform, Institute of Integrative and Systems Biology, Université Laval (Quebec, Canada) for GbS library preparation. DNA was digested with a combination of restriction enzymes PstI/NsiI/MspI, ligated with barcoded adapter, and pooled to equimolarity. The pooled library was amplified by PCR and sequenced on a Illumina NovaSeq6000 S4 with paired-end reads of 150 bp at the Génome Québec Innovation Centre, (Montreal, Canada).

### Variant calling

DNA sequence variant calling was done with the Fast-GBS v2.0 pipeline^79^: Illumina raw reads were demultiplexed with Sabre 1.0^80^ and trimmed with Cutadapt 1.18^81^ to remove the adaptors. Trimmed reads shorter than 50 bp were discarded. Reads were aligned against the Dacoc 1.0 genome (Hung et al., unpublished) and the Daoli 0.1 genome using BWA-MEM 0.7.17^82^. The SAM alignment files were converted to BAM format and indexed using SAMtools 1.9^83^. Variant calling was performed in Platypus^84^ and variants were filtered with proportion of missing data of 0.2 and minimum allele frequency (MAF) of 0.01 using VCFtools 0.1.16^85^. Missing genotype was imputed using Beagle 5.2. Finally, linkage equilibrium among SNPs was detected using BCFtools 1.9^83^, and one SNP was removed from all SNP pairs with *r*^2^ > 0.5 in a genomic window of 5 Kbp.

### Environmental heterogeneity characterisation

Environmental data were obtained from different sources (34 variables in total, Supplementary Table 12) and represented different measurers of temperature, precipitation, their seasonality, soil, elevation, and vegetation. We calculated a correlation matrix across the sampling localities and highly inter-correlated variables (pairwise correlation coefficient| > 0.7) were detected. For each inter-correlated variable pair, the one variable with the largest mean absolute correlation across all variables was removed.

### Population genetic structure and identification of putatively adaptive loci

Population genetic structure was assessed with sparse non-negative matrix factorisation (sNMF) to estimate the number of discrete genetic clusters (K)^86^. The sNMF was run for 10 repetitions for each value of K from 1 to 15 with a maximum iteration of 200. The optimal K was selected based on the lowest cross-entropy value from the sNMF run, or where the value began to plateau. Admixture plots were drawn for K = {2, 4, 8, optimal K}. Population structure-based outlier analysis was also conducted with sNMF, in which outlier SNPs that are significantly differentiated among populations, based on estimated F_ST_ values from the ancestry coefficients obtained from sNMF^87^, were obtained and mapped on the 10 putative chromosomes for *D. cochinchinensis* or the 16 longest scaffolds for *D. oliveri* in a Manhattan plot.

We used latent factor mixed modelling (LFMM) to test for significant associations between environmental variables and SNP allele frequencies. The optimal K obtained from the sNMF was used in LFMM to correct for the neutral genetic structure. LFMM was run for 3 repetitions with a maximum iteration of 1,000 and 500 burn-ins. Z-scores were obtained for all repetitions for each environmental variable, and then the median was taken for each SNP. Next, the genomic inflation factor λ, defined as the observed median of Z-scores divided by the expected median of the chi-squared distribution for each environmental association^88^, was calculated to calibrate for *P-*values:

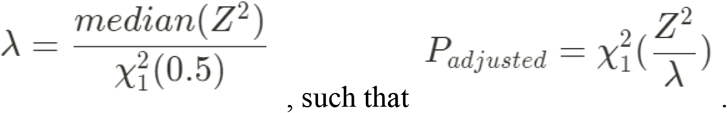

The calibration was then inspected on a histogram of *P-*values for each environmental association. Finally, multiple testing was corrected with the Benjamini and Hochberg method to obtain *Q*-values.

The sNMF and LFMM calculations were performed in R 4.1.0 using the packages LEA 3.4.0^89^.

### Gradient forest modelling

For all predictions in gradient forest models, resampling was necessary because not all environmental raster layers had the same resolution and extent. They were all cropped to the latest-updated modelled and expert-validated species distribution^12^ and reprojected to the WorldClim bioclimatic rasters, as they have the highest resolution, using bilinear interpolation or nearest neighbour method for continuous and categorical variables respectively.

To correct for the genetic structure, spatial variables were generated using the principal coordinates of neighbour matrices (PCNM) approach^90^. Only half of the positive PCNM values were kept. Gradient forest model was used to predict and rank the importance of environmental variables in genomic variation, as its machine learning algorithm worked best with minimal prior and confounding variables. Putatively neutral SNPs and putatively adaptive SNPs were used as the response variables and all the filtered environment variables and PCNM variables were used as the predictor variables in the gradient forest model for 500 regression trees. The maximum number of splits to evaluate was determined as follows:

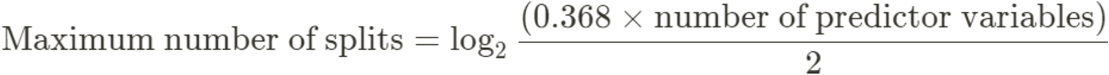

The turnovers of allelic frequencies were then projected spatially across the latest-updated predicted species distribution ranges^12^ using the fitted gradient forest model and the environmental values across the range. Principal component analysis (PCA) was used to summarise the genomic variation across the distribution and the first three principal components (PC1, PC2, and PC3) were used for visualisation of genomic variation across the range.

The PCAs of turnovers of allelic frequencies between adaptive SNPs and neutral SNPs were compared using the Procrustes rotation, and its residuals were used to map where adaptive genomic variation deviates from neutral variation.

### Prediction of genomic vulnerability

Future climate projections were obtained from five general circulation models (GCM) (MIROC6, BCC-CSM2-MR, IPSL-CM6A-LR, CNRM-ESM2-1, MRI-ESM2-0) participating in the World Climate Research Programme Coupled Model Intercomparison Project 6 (WCRP CMIP6) for four shared socio-economic pathways (SSPs) (126, 245, 370, and 585) over four 20-year periods (2021–2040, 2041–2060, 2061–2080, 2081–2100). The gradient forest model was used to predict patterns of genetic variation and local adaption under future environmental scenarios. The allelic frequency turnover function was fitted on the future landscape and the genomic offset, defined as the required genomic change in a set of putatively adaptive loci to adapt to a future environment^91^, was calculated in a grid-by-grid basis using the following equation for Euclidean distance, where *p* is the number of environmental (predictor) variables:

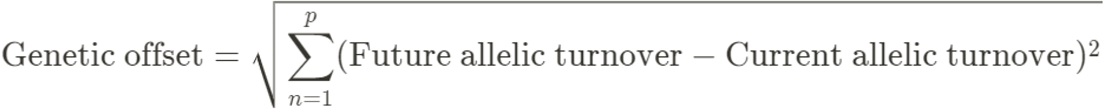

The genetic offset was then scaled across all SSPs and time periods.

### Prediction of genomic similarity between current germplasm source and future restoration site

It is of practical interest to a range of forestry stake-holders to predict if a current germplasm source is a good match for future restoration sites, or where to source suitable germplasm for a proposed restoration site. We developed an interactive web application based on R Shiny and hosted the application on the shinyapps.io server. *seedeR* v 1.0 is open source and freely available from https://trainingidn.shinyapps.io/seeder/. The analysis workflow consists of the selection of species of interest, time period and future climate scenario, and the restoration site’s geographical coordinates (Supplementary Figure 11).

The application maps the predicted turnover of allelic frequencies at a hypothetical future restoration site onto the current landscape on a grid-by-grid basis, with the genetic offset calculated as described above. After scaling, the values are reversed on a 0-1 scale to represent the genomic similarity between the current germplasm source and future restoration site.

## Data availability

The research materials supporting this publication, including genomic assemblies, raw reads, and annotations, can be publicly accessed either in the Supplementary Information or in NCBI GenBank under the BioProjects PRJNA841235 and PRJNA841689.

## Author contributions

T.H.H.: designed the study, processed the samples, conducted the Oxford Nanopore sequencing, conceived and conducted the bioinformatic analyses, drafted the manuscript, and secured funding for the project;

T.S: collected the samples, revised the manuscript, and secured funding for the project;

B.T.: collected the samples, revised the manuscript, and secured funding for the project;

V.C.: collected the samples, and revised the manuscript;

I.T.: collected the samples, revised the manuscript, and secured funding for the project;

C.P.: collected the samples, and revised the manuscript;

S.B.: collected the samples, and revised the manuscript;

I.H.: collected the samples, and revised the manuscript;

H.G.: provided expertise and materials for species distribution models, and revised the manuscript;

R.J.: revised the manuscript, and secured funding for the project;

D.H.B.: supervised the study, revised the manuscript, and secured funding for the project;

J.J.M.: designed and supervised the study, revised the manuscript, and secured funding for the project.

## Acknowledgements

The genomic work was supported by funding to T.H.H. from the Biotechnology and Biological Sciences Research Council (grant number BB/M011224/1) and to T.H.H., J.J.M. from the Google Cloud Academic Grant. The sampling work was supported by funding to T.S., B.T., I.T., R.J., D.H.B., J.J.M from the UK Darwin Initiative (ref. 25-023). The work of H.G. and R.J. was supported by the CGIAR Fund Donors (https://www.cgiar.org/funder) through the CGIAR Research Programme on Forests, Trees and Agroforestry. T.H.H. wishes to thank Andrew Eckert and Stephen Harris as the examiners of his doctoral thesis, who provided very constructive feedback that improved this paper.

