## Supplementary Information for "Range-wide differential adaptation and genomic vulnerability in critically endangered Asian rosewoods"

Supplementary Table 1.

Assembly statistics of the *D. cochinchinensis* genome (Dacoc_1.4).

| | |
|---|---|
| Assembly identifier | Dacoc_1.4 |
| Species | *Dalbergia cochinchinensis* Pierre ex Laness |
| NCBI taxonomy ID | 106130 |

| | ***Raw reads*** | ***Hi-C reads*** | ***Contigs*** | ***Primary haplotigs*** | ***Scaffolds*** |
|---|---|---|---|---|---|
| Total size (Mbp) | | 54,973.60 | 1,353.09 | 621.01 | 621.01 |
| Number of sequences | | 366,490,675 | 6,443 | 1,238 | 512 (?) |
| Average length of sequences | | 150 | 210,008.8 | 1,003,247.1 | 1,212,910.1 |
| N50 | | 150 | 1,345,847 | 21,352,153 | 59,971,594 |

Supplementary Table 2.

Statistics of Hi-C library prepared with the restriction enzyme DpnII and proximity-ligated.

| | |
|---|---|
| Total read pairs (RPs) | 366,490,675 |
| RP length | 150 bp |
| High quality (HQ) RPs | 25.37% |
| RPs > 10 Kb apart | 6.82% |
| RPs > 10 Kb apart (CTGs > 10 Kb) | 10.37% |
| Intercontig RPs | 33.22% |
| Intercontig hig-quality (HQ) RPs | 6.31% |
| Same strand RPs | 7.22% |
| Split reads | 13.53% |
| Duplicate reads (extrapolated) | 8.72% |
| Genome scaffolding sufficiency | |
| Same-strand high-quality* (HQ) read pairs (RPs) | 5.03% |
| Informative RPs** | 5.21% |

Supplementary Table 3.

Assembly statistics of the *D. oliveri* genome (Daoli_0.3).

| | |
|---|---|
| Assembly identifier | Daoli_0.3 |
| Species | *Dalbergia oliveri* Gamble ex Prain |
| NCBI taxonomy ID | 1030150 |

| | ***Raw reads*** | ***Pore-C reads*** | ***Contigs*** | ***Primary haplotigs*** | ***Scaffolds*** |
|---|---|---|---|---|---|
| Total size (Mbp) | 15,127.06 | 13,462.40 | 814.69 | 687.92 | 689.25 |
| Number of sequences | 959,359 | 3,098,972 | 3,249 | 2,239 | 2,977 |
| Average length of sequences | 15,767.9 | 4,344.2 | 250,750 | 307,244.3 | 231,525 |
| N50 | 44,832 | 5,789 | 474,015 | 623,924 | 38,426,337 |

Supplementary Figure 1.

Snailplot of the genome assembly (a) Dacoc_1.4 and (b) Daoli_0.3 showing scaffold statistics, BUSCO statistics, and assembly base composition.

(a)

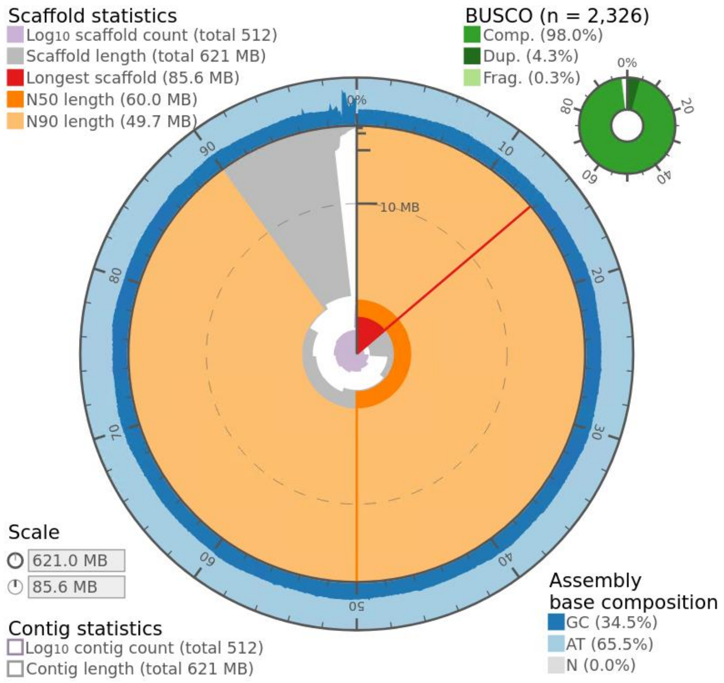


(b)

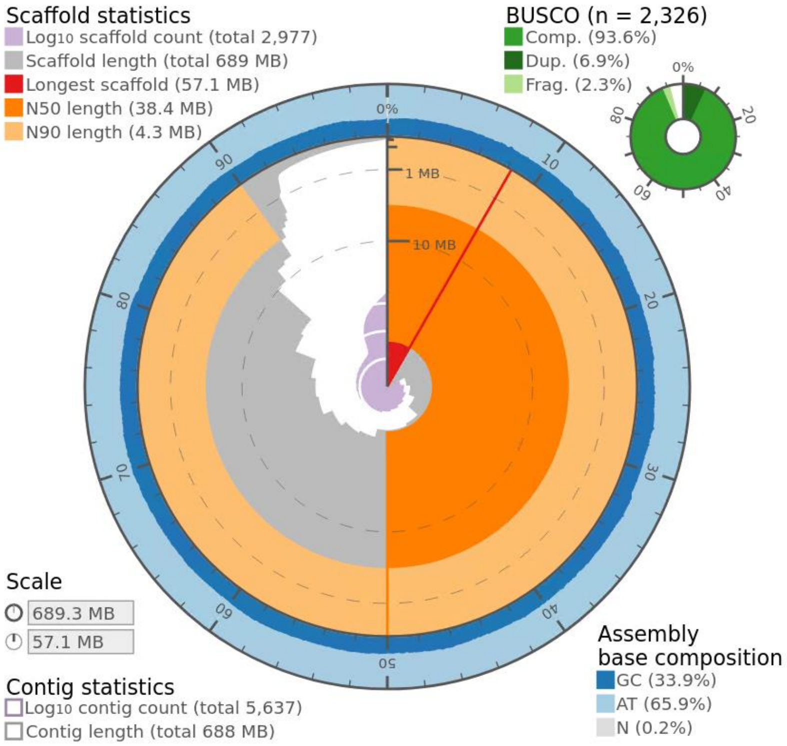

Supplementary Table 4.

Composition and percentage of the repeat families in the final genome of Dacoc_1.4 identified with RepeatModeler and LTRharvest.

| | **Number of elements** | **Total length (bp)** | **Percentage of sequence** |
|---|---|---|---|
| **Dacoc_1.2** | | 621009970 | |
| **Bases masked** | | 402405433 | 64.80 |
| **Retroelements** | 405732 | 293464285 | 47.26 |
| SINEs | 0 | 0 | 0.00 |
| Penelope | 0 | 0 | 0.00 |
| LINEs | 9518 | 3856482 | 0.62 |
| CRE/SLACS | 0 | 0 | 0.00 |
| L2/CR1/Rex | 0 | 0 | 0.00 |
| R1/LOA/Jockey | 0 | 0 | 0.00 |
| R2/R4/NeSL | 0 | 0 | 0.00 |
| RTE/Bov-B | 3633 | 803221 | 0.13 |
| L1/CIN4 | 5885 | 3053261 | 0.49 |
| LTR elements | 396214 | 289607803 | 46.63 |
| BEL/Pao | 0 | 0 | 0.00 |
| Ty1/Copia | 209546 | 94712214 | 15.25 |
| Gypsy/DIRS1 | 176990 | 189459322 | 30.51 |
| Retroviral | 0 | 0 | 0.00 |
| **DNA transposons** | 11462 | 6635025 | 1.07 |
| hobo-Activator | 1830 | 871791 | 0.14 |
| Tc1-IS630-Pogo | 0 | 0 | 0.00 |
| En-Spm | 0 | 0 | 0.00 |
| MuDR-IS905 | 0 | 0 | 0.00 |
| PiggyBac | 0 | 0 | 0.00 |
| Tourist/Harbinger | 3272 | 1401349 | 0.23 |
| Other (Mirage, P-element, Transib) | 0 | 0 | 0.00 |
| **Rolling-circles** | 3613 | 2108774 | 0.34 |
| **Unclassified** | 359305 | 90585676 | 14.59 |
| Total interspersed repeats | | 390684986 | 62.91 |
| Simple repeats | 131568 | 8474809 | 1.36 |
| Low complexity | 23616 | 1136864 | 0.18 |

Supplementary Table 5.

Composition and percentage of the repeat families in the final genome of Daoli_0.3 identified with RepeatModeler and LTRharvest.

| | **Number of elements** | **Total length (bp)** | **Percentage of sequence** |
|---|---|---|---|
| **Retroelements** | 455479 | 339092238 | 49.2 |
| SINEs | 0 | 0 | 0 |
| Penelope | 0 | 0 | 0 |
| LINEs | 10758 | 4445290 | 0.64 |
| CRE/SLACS | 0 | 0 | 0 |
| L2/CR1/Rex | 0 | 0 | 0 |
| R1/LOA/Jockey | 0 | 0 | 0 |
| R2/R4/NeSL | 0 | 0 | 0 |
| RTE/Bov-B | 3783 | 973287 | 0.14 |
| L1/CIN4 | 6975 | 3472003 | 0.5 |
| LTR elements | 444721 | 334646948 | 48.55 |
| BEL/Pao | 0 | 0 | 0 |
| Ty1/Copia | 229457 | 108551031 | 15.75 |
| Gypsy/DIRS1 | 200968 | 220263371 | 31.96 |
| Retroviral | 181 | 91342 | 0.01 |
| **DNA transposons** | 16939 | 12338687 | 1.89 |
| hobo-Activator | 2978 | 1273955 | 0.18 |
| Tc1-IS630-Pogo | 1440 | 179489 | 0.03 |
| En-Spm | 0 | 0 | 0 |
| MuDR-IS905 | 0 | 0 | 0 |
| PiggyBac | 0 | 0 | 0 |
| Tourist/Harbinger | 2277 | 1185567 | 0.17 |
| Other (Mirage, P-element, Transib) | 0 | 0 | 0 |
| **Rolling-circles** | 5146 | 3333791 | 0.48 |
| **Unclassified** | 376428 | 91174602 | 13.23 |
| Total interspersed repeats | 1757530 | 442605527 | 64.22 |
| Simple repeats | 143462 | 5735706 | 0.83 |
| Low complexity | 25174 | 1218960 | 0.18 |

Supplementary Table 6.

Mapping statistics of RNA-seq raw reads from a previous experiment using pooled sample_from leaves, stems, and roots of *D. cochinchinensis* and *D. oliveri* on the Dacoc_1.4 and Daoli_0.3 genomes respectively.

| | Dacoc_1.4 | Daoli_0.3 |
|---|---|---|
| Number of input reads | 135,924,672 | 132,696,766 |
| Average input read length | 299 | 299 |
| | ***Unique reads*** | |
| Number of uniquely mapped reads | 115,815,104 (85.21%) | 122,215,722 (92.10%) |
| Average mapped length | 294.91 | 296.69 |
| Number of splices<br>GT/AG<br>GC/AG<br>AT/AC<br>Non-canonical | 86,392,015<br>84,315,981<br>956,359<br>47,749<br>1,071,926 | 96,847,851<br>95,280,260<br>1,148,136<br>44,025<br>375,430 |
| Mismatch rate per base (%) | 0.86 | 2.87 |
| Deletion rate per base (%) | 0.05 | 0.03 |
| Deletion average length | 2.95 | 2.87 |
| Insertion rate per base (%) | 0.11 | 0.03 |
| Insertion average length | 1.62 | 2.50 |
| | ***Multi-mapping reads*** | |
| Number of reads mapped to multiple loci | 11,182,606 (8.23%) | 4,875,679 (3.67%) |
| Number of reads mapped to too many loci | 27,541 (0.02%) | 230,932 (0.17%) |
| | ***Unmapped reads*** | |
| Number of reads with too many mismatches | 0 (0.00%) | 0 (0.00%) |
| Number of reads too short | 8,890,737 (6.54%) | 4,954,750 (3.73%) |
| Number of other reads | 8,684 (0.01%) | 419,683 (0.32%) |
| | ***Chimeric reads*** | |
| Number of chimeric reads | 0 (0.00%) | 0 (0.00%) |

Supplementary Figure 2.

Cumulative frequency of gene models of **(a)** Dacoc_1.4 and **(b)** Daoli_0.3 against annotation edit distance (AED).

(a)

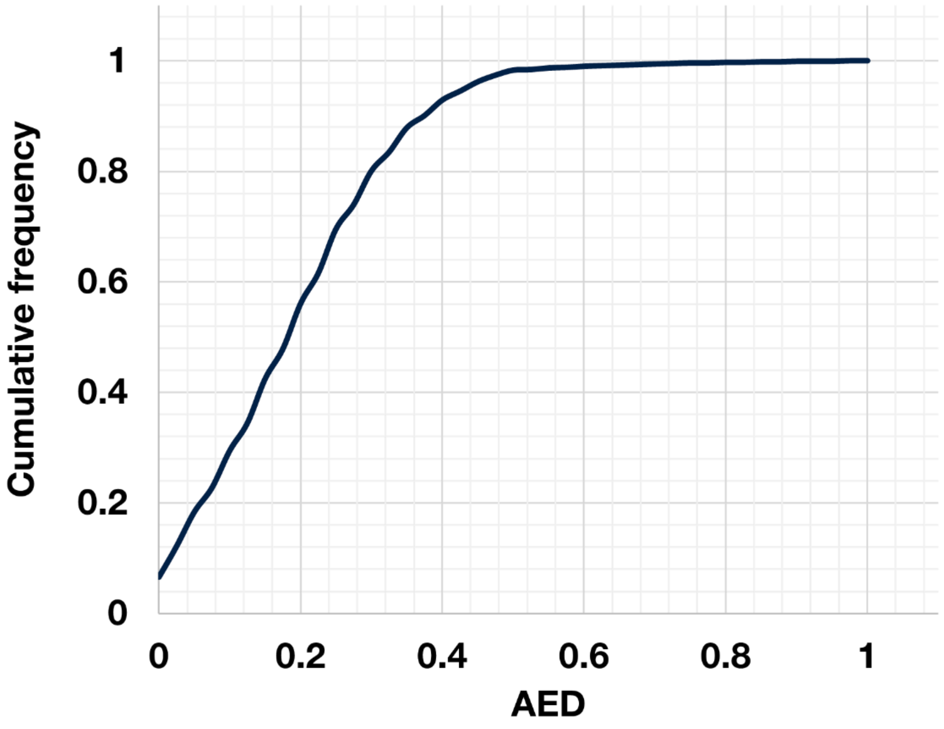


(b)

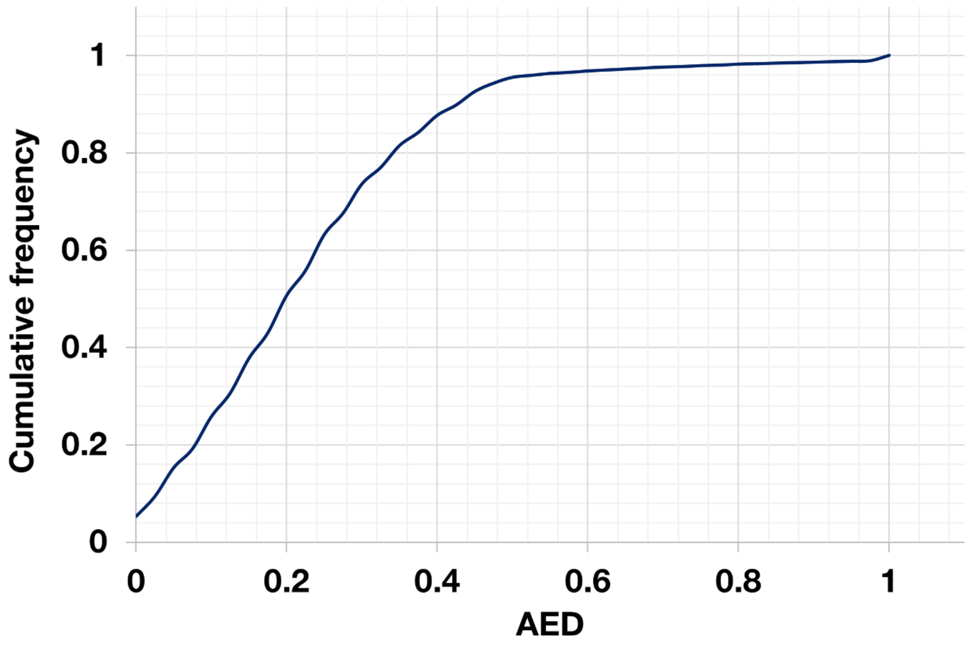

Supplementary Table 7.

Basic statistics of the ecological genomic study in *D. cochinchinensis* and *D. oliveri*.

| | ***D. cochinchinensis*** | ***D. oliveri*** |
|---|---|---|
| # Samples | 435 | 331 |
| # Localities | 35 | 28 |
| # SNPs after variant calling | 1,832,629 | 3,377,855 |
| # SNPs after filtering for missing data and MAF | 246,225 | 250,954 |
| # SNPs after pruning for linkage disequilibrium | 180,944 | 193,724 |
| # Adaptive SNPs predicted by sNMF | 50,951 | 54,154 |
| # Adaptive SNPs predicted by LFMM | 20,373 | 6,953 |

Supplementary Figure 3.

Cross-entropy criterion as a function of the number of factors in the sNMF analysis for **(a)** *D. cochinchinensis* and **(b)** *D. oliveri*.

**(a)**

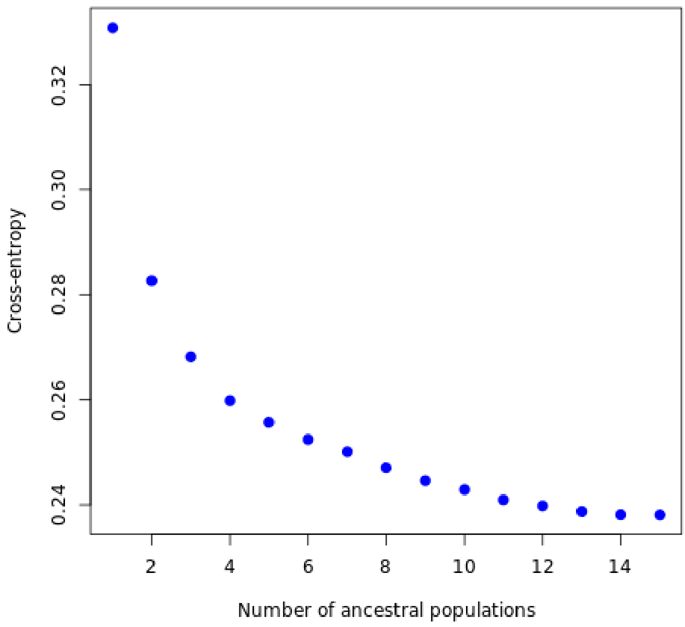


**(b)**

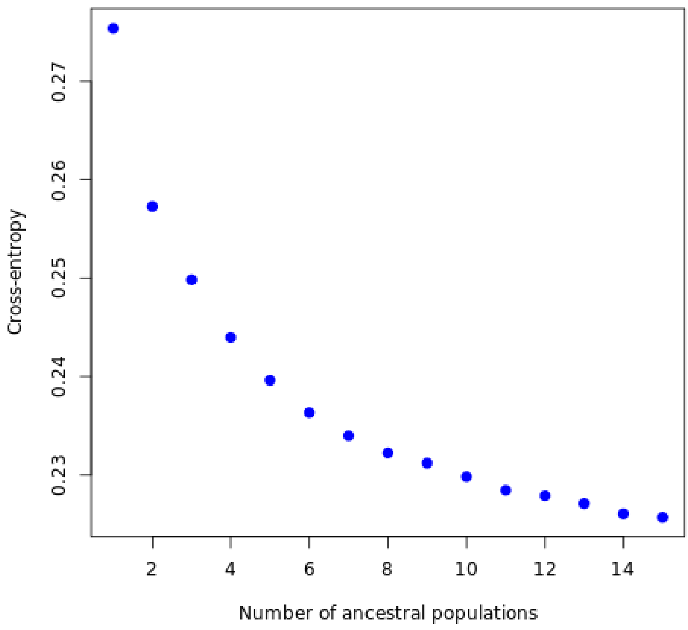

Supplementary Figure 4.

Admixture results of sNMF analysis showing hierarchical population substructure of **(a)** *D. cochinchinensis* and **(b)** *D. oliveri*.

**(a)**

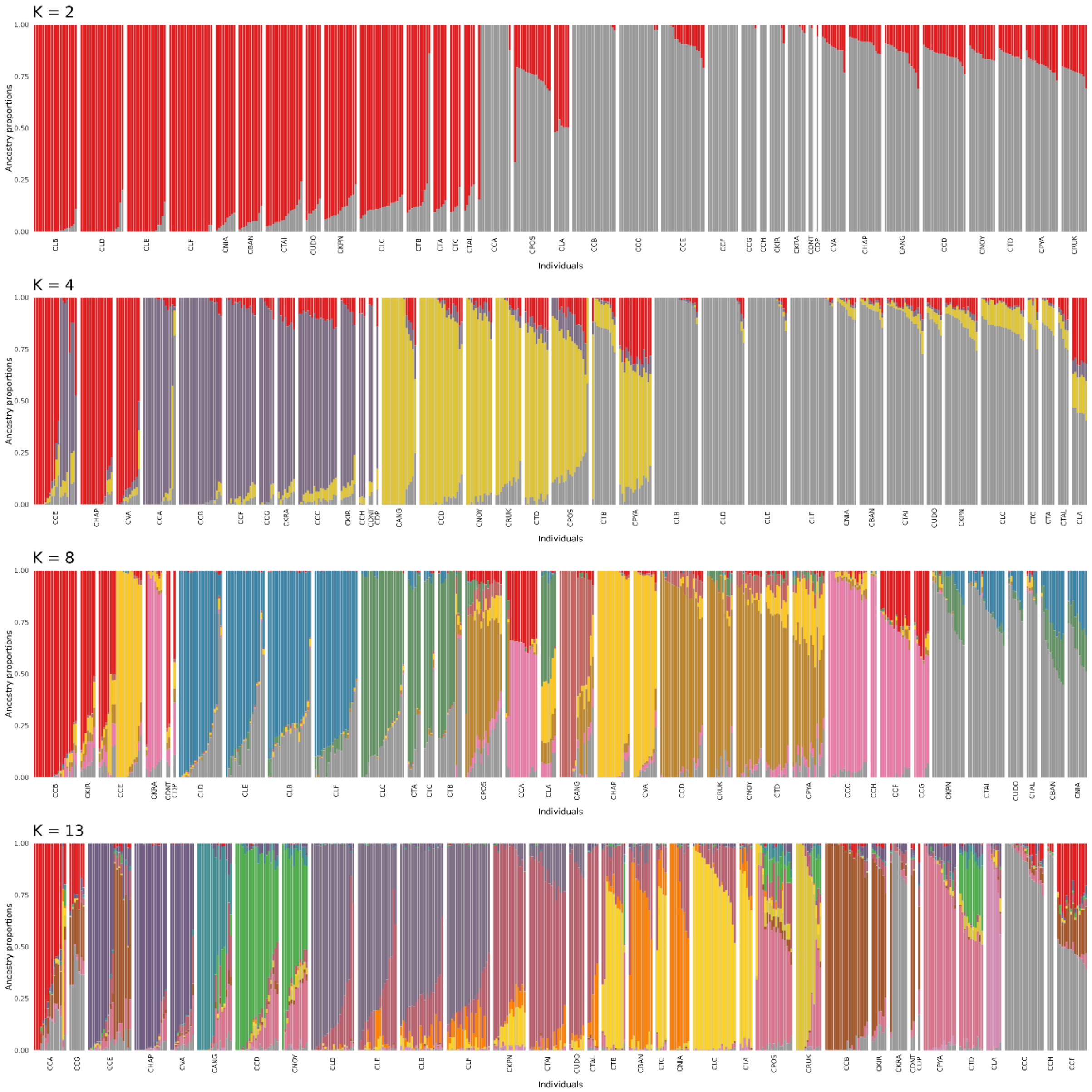

**(b)**

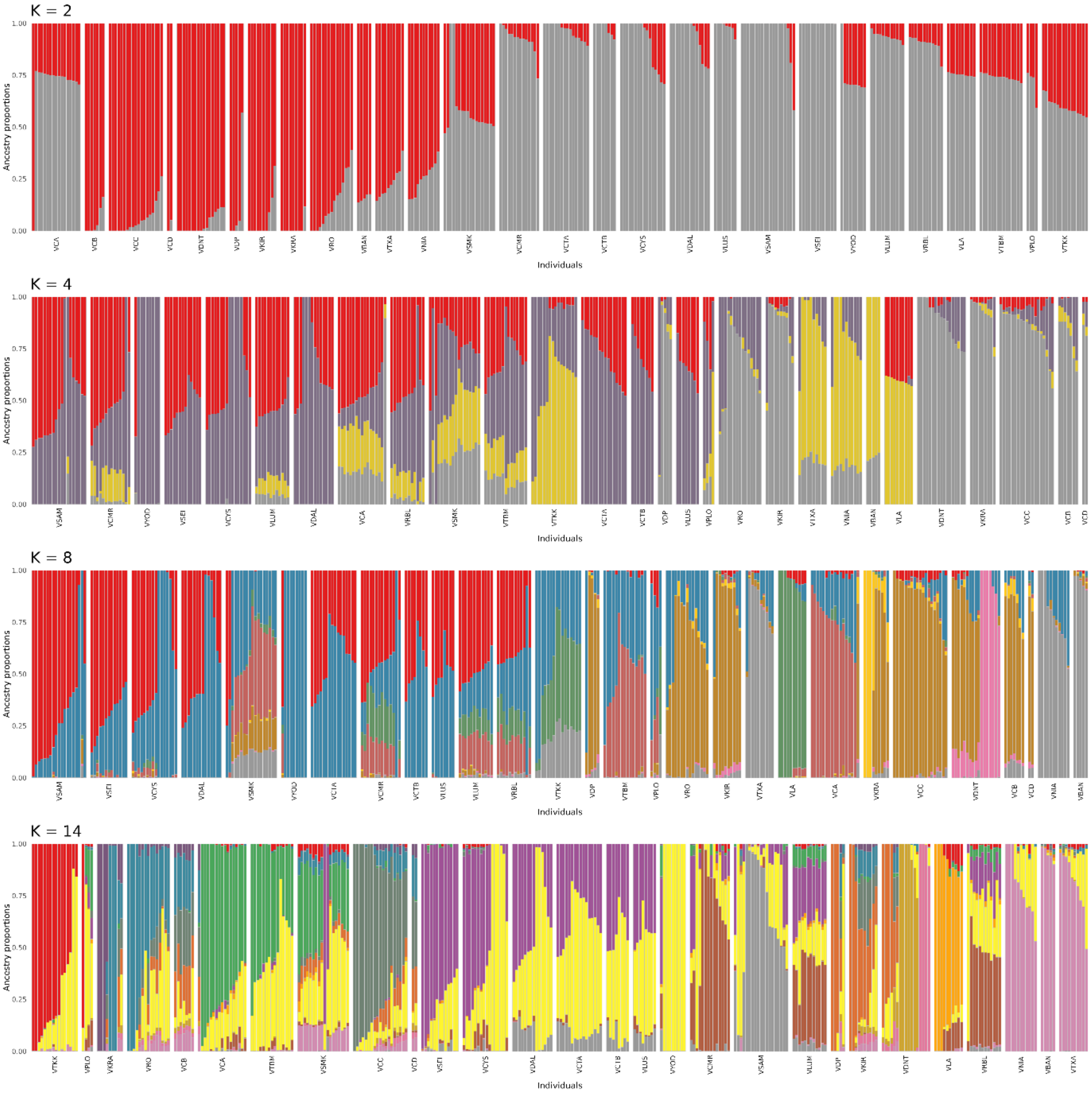
K = 2
K = 4
K = 8
K = 14
Ancestry proportions
Individuals
0.00
0.25
0.50
0.75
1.00

Supplementary Figure 5.

Admixture results of sNMF analysis showing hierarchical population substructure of **(a)** *D. cochinchinensis* and **(b)** *D. oliveri*.

**(a)**

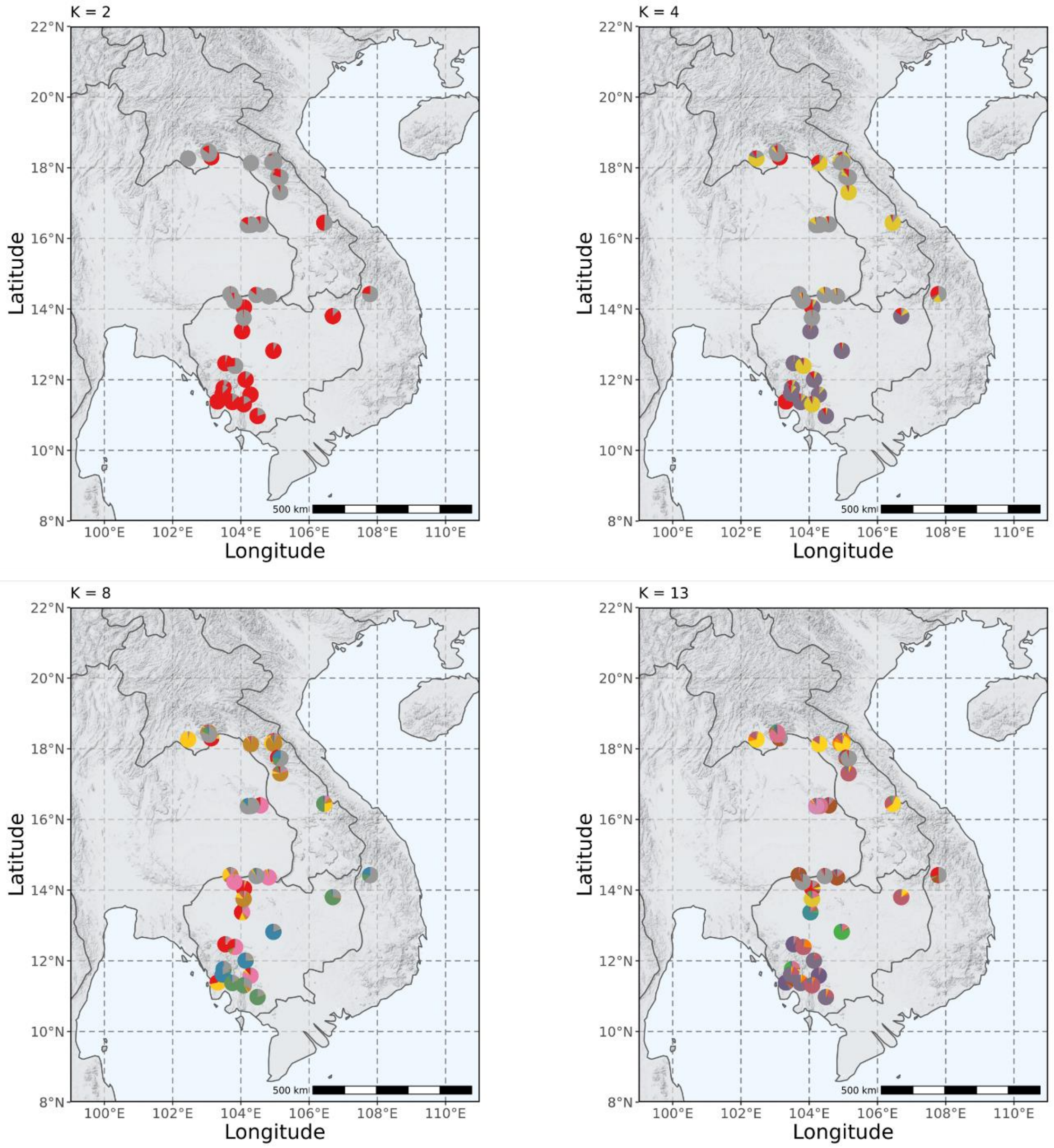

**(b)**

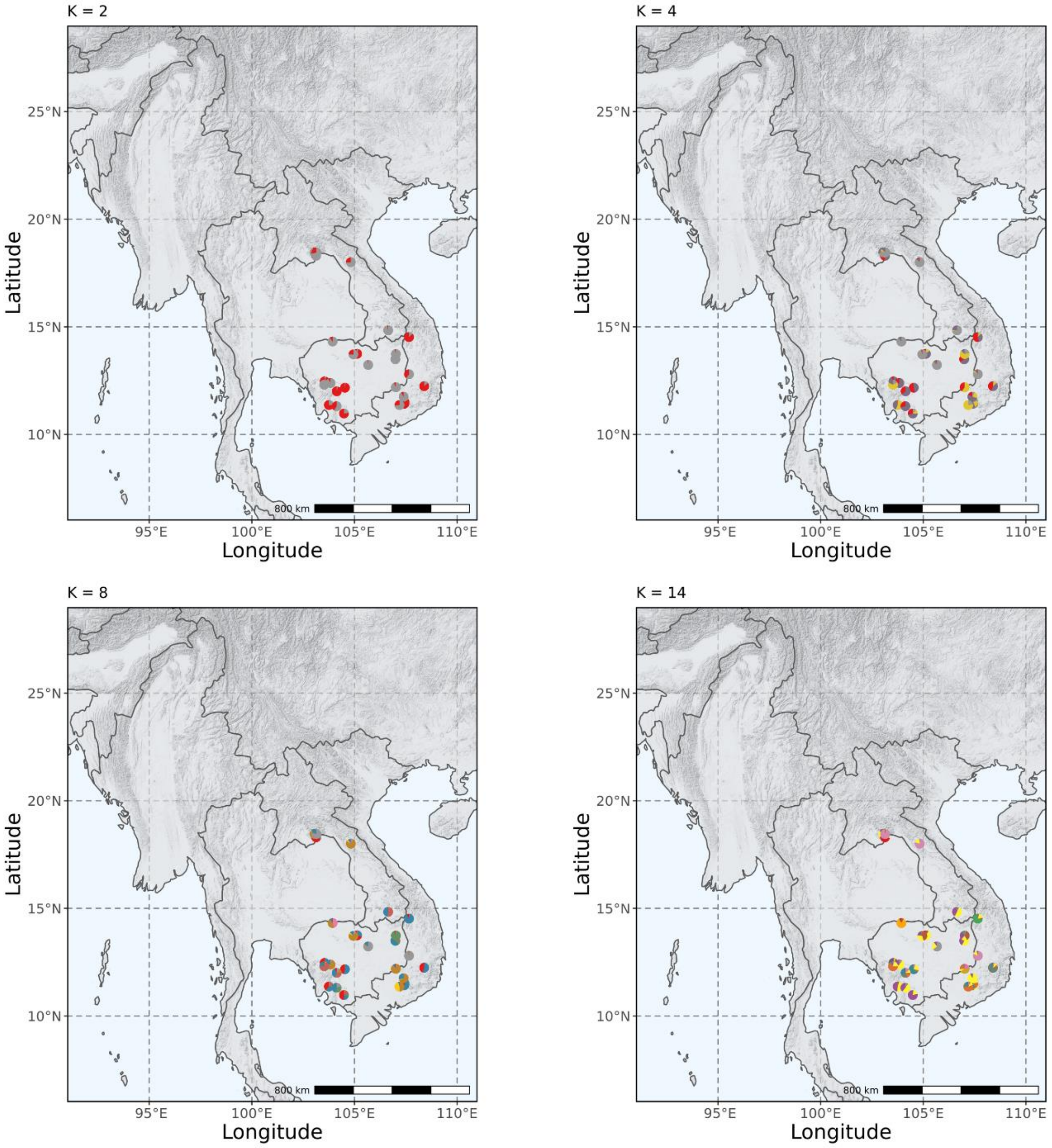
K = 2
K = 4
K = 8
K = 14
25°N
20°N
15°N
10°N
Latitude
95°E
100°E
105°E
110°E
Longitude
800 km

Supplementary Table 8.

Genomic inflation factor λ for the gene-environment associations of **(a)** *D. cochinchinensis* and **(b)** *D. oliveri*

| env | lambda |
|---|---|
| evapotrans | 0.071053 |
| s_AWCh1_sl2 | 0.1012 |
| s_AWCh1_sl5 | 0.097883 |
| s_CLYPPT_sl2 | 0.0773 |
| s_ORCDRC_sl5 | 0.103751 |
| s_PHIHOX_sl5 | 0.11886 |
| s_SLTPPT_sl2 | 0.117395 |
| wc2.1_30s_bio_13 | 0.166612 |
| wc2.1_30s_bio_17 | 0.247199 |
| wc2.1_30s_bio_19 | 0.136046 |
| wc2.1_30s_bio_2 | 0.186022 |
| wc2.1_30s_bio_3 | 0.144369 |
| wc2.1_30s_bio_8 | 0.088284 |
| **mean** | **0.127383** |
| **sd** | **0.049323** |

| env | lambda |
|---|---|
| evapotrans | 0.038403 |
| npp | 0.046587 |
| s_AWCh1_sl2 | 0.037444 |
| s_AWCh1_sl5 | 0.057747 |
| s_ORCDRC_sl2 | 0.05131 |
| s_PHIHOX_sl2 | 0.05708 |
| s_SLTPPT_sl5 | 0.050496 |
| s_SNDPPT_sl5 | 0.032456 |
| wc2.1_30s_bio_14 | 0.070037 |
| wc2.1_30s_bio_16 | 0.079196 |
| wc2.1_30s_bio_2 | 0.080899 |
| wc2.1_30s_bio_3 | 0.069888 |
| **mean** | **0.055962** |
| **sd** | **0.016241** |

Supplementary Figure 6.

Manhattan plot of adaptive SNPs predicted by LFMM for **(a)** *D. cochinchinensis* and **(b)** *D. oliveri*. Blue dots are adaptive SNPs for at least one environmental variable ( | *Z*-value | > 2 & *Q*-value < 0.01), light-blue dots are neutral SNPs.

**(a)**

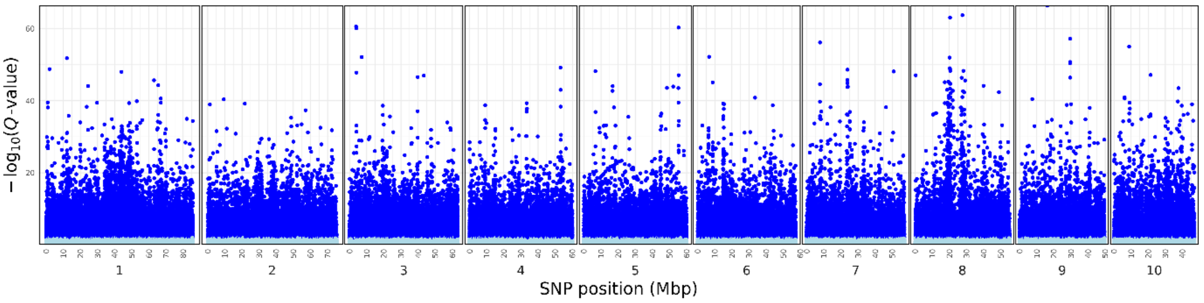


**(b)**

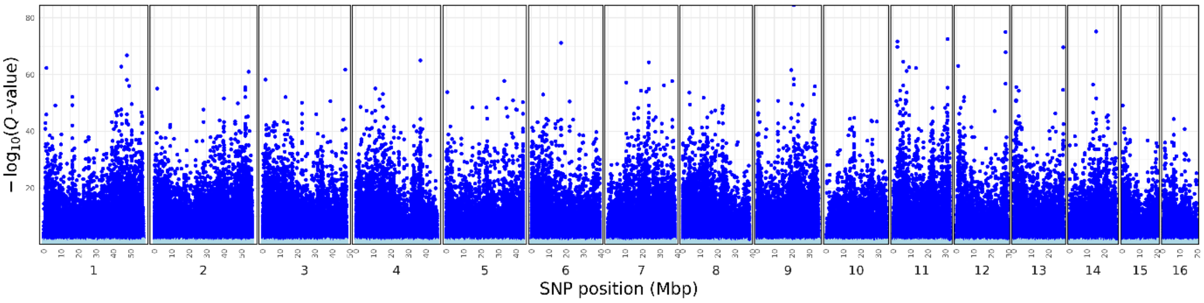

Supplementary Figure 7.

Frequency histogram of adjusted *P*-values for each environmental association of the LFMM analysis in **(a)** *D. cochinchinensis* and **(b)** *D. oliveri*.

**(a)**

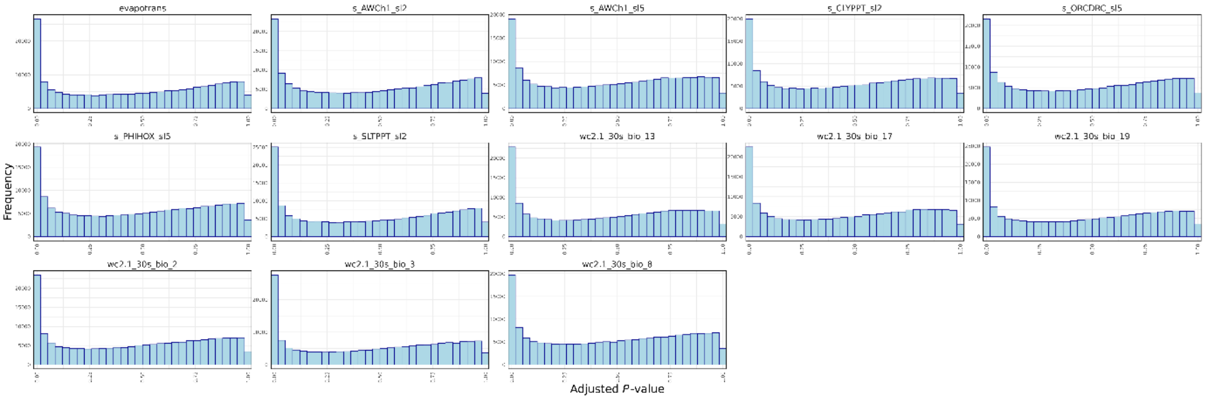


**(b)**

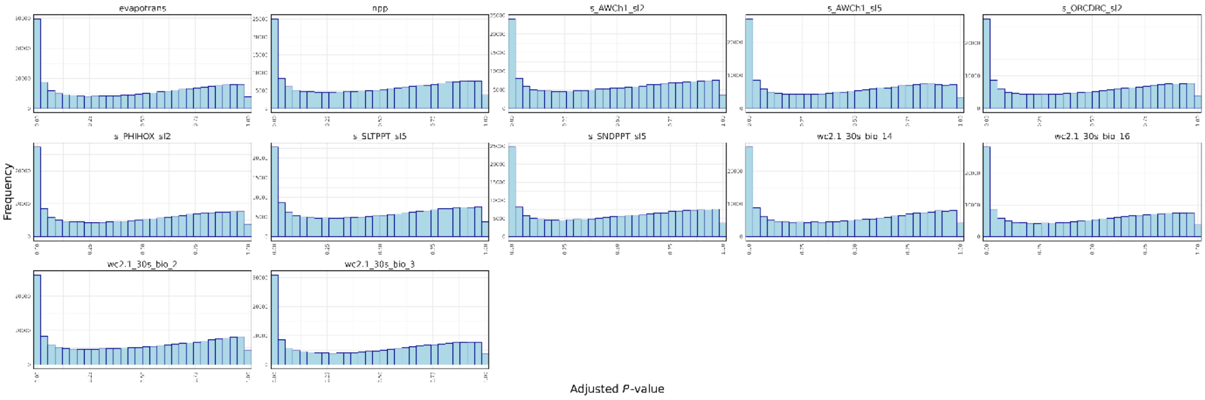

Supplementary Table 9.

Absolute *Z*-values and *Q*-values of putatively adaptive SNPs predicted by LFMM, only showing the significant SNPs for at least one environmental variable ( | *Z*-value | > 2 & *Q*-value < 0.01).

*In a separate spreadsheet*

Supplementary Figure 8.

Accuracy and $R^2$-weighted importance for the environmental predictor variables which explained the neutral genomic variation (non-adaptive SNPs) for **(a)** *D. cochinchinensis* and **(b)** *D. oliveri* predicted by the GF model.

**(a)**

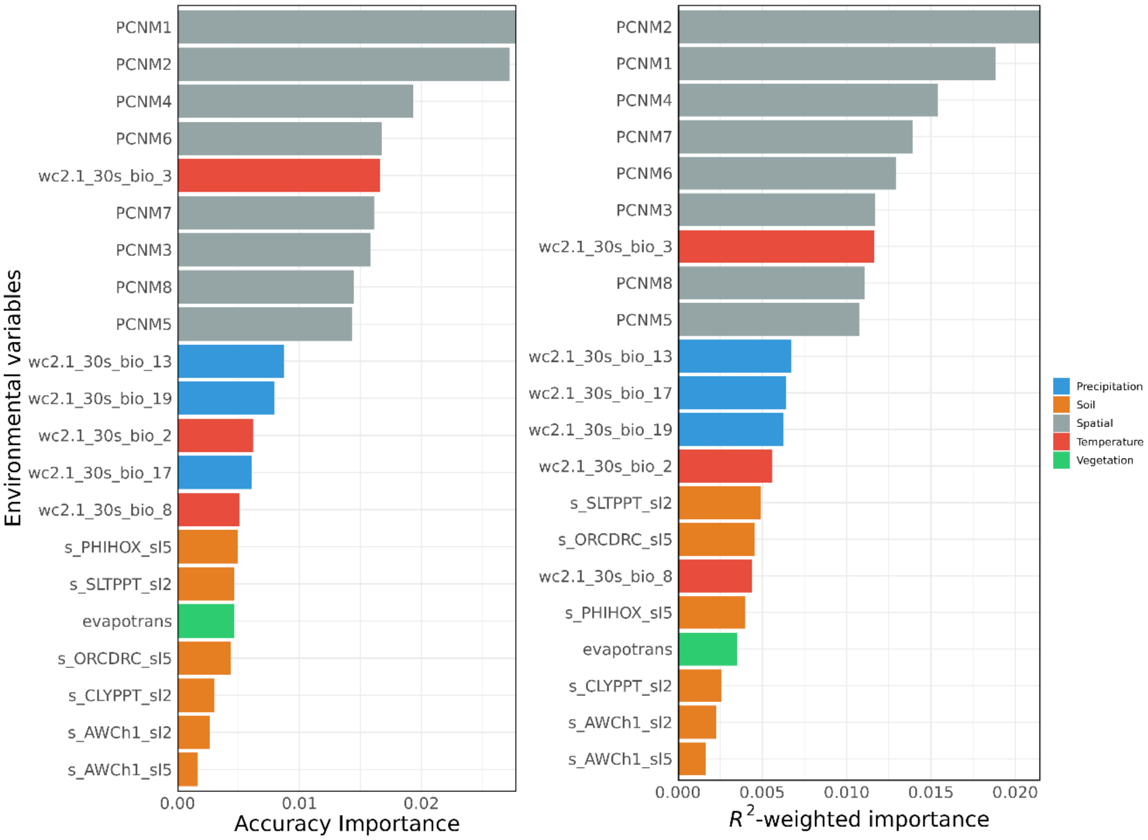

**(b)**

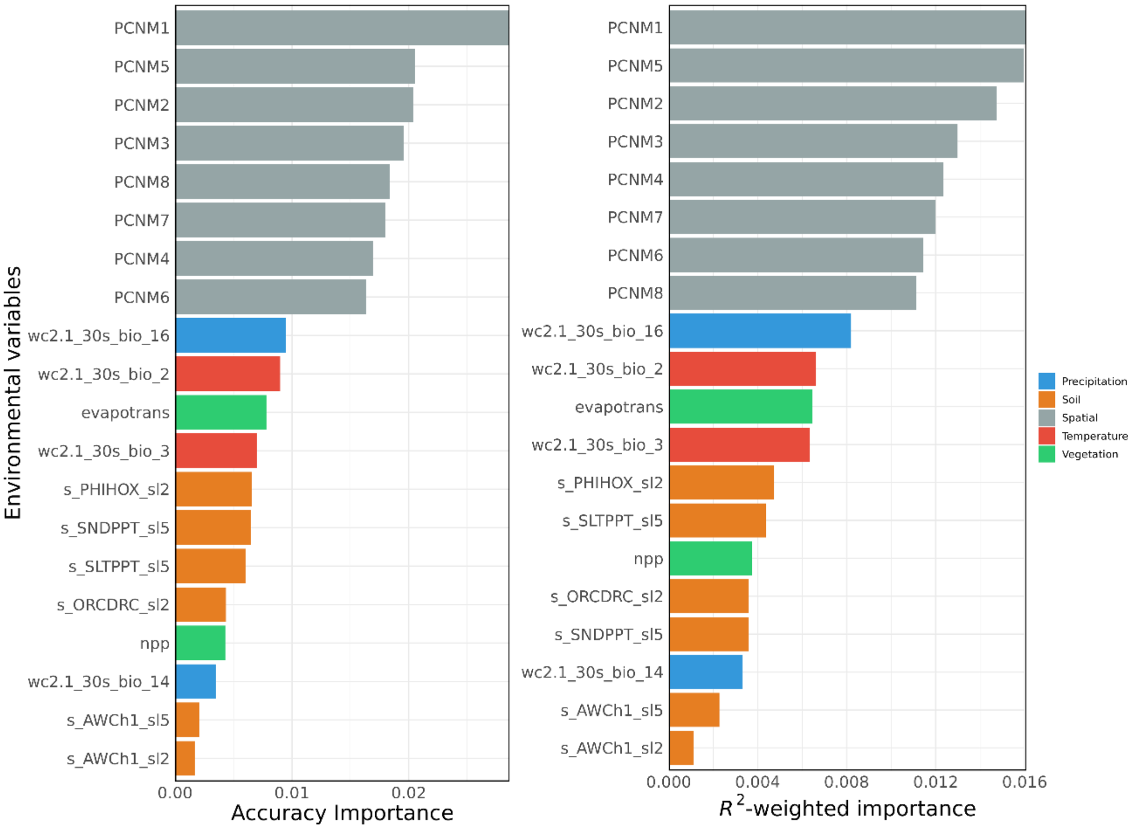
Environmental variables
PCNM1
PCNM5
PCNM2
PCNM3
PCNM8
PCNM7
PCNM4
PCNM6
wc2.1_30s_bio_16
wc2.1_30s_bio_2
evapotrans
wc2.1_30s_bio_3
s_PHIHOX_sl2
s_SNDPPT_sl5
s_SLTPPT_sl5
s_ORCDRC_sl2
npp
wc2.1_30s_bio_14
s_AWCh1_sl5
s_AWCh1_sl2
0.00
0.01
0.02
Accuracy Importance
PCNM1
PCNM5
PCNM2
PCNM3
PCNM4
PCNM7
PCNM6
PCNM8
wc2.1_30s_bio_16
wc2.1_30s_bio_2
evapotrans
wc2.1_30s_bio_3
s_PHIHOX_sl2
s_SLTPPT_sl5
npp
s_ORCDRC_sl2
s_SNDPPT_sl5
wc2.1_30s_bio_14
s_AWCh1_sl5
s_AWCh1_sl2
0.000
0.004
0.008
0.012
0.016
$R^2$-weighted importance
Precipitation
Soil
Spatial
Temperature
Vegetation

Supplementary Figure 9.

Allelic turnover functions of all and adaptive SNPs explaining the cumulative importance of each environmental predictor variable in **(a)** *D. cochinchinensis* and **(b)** *D. oliveri*.

**(a)**

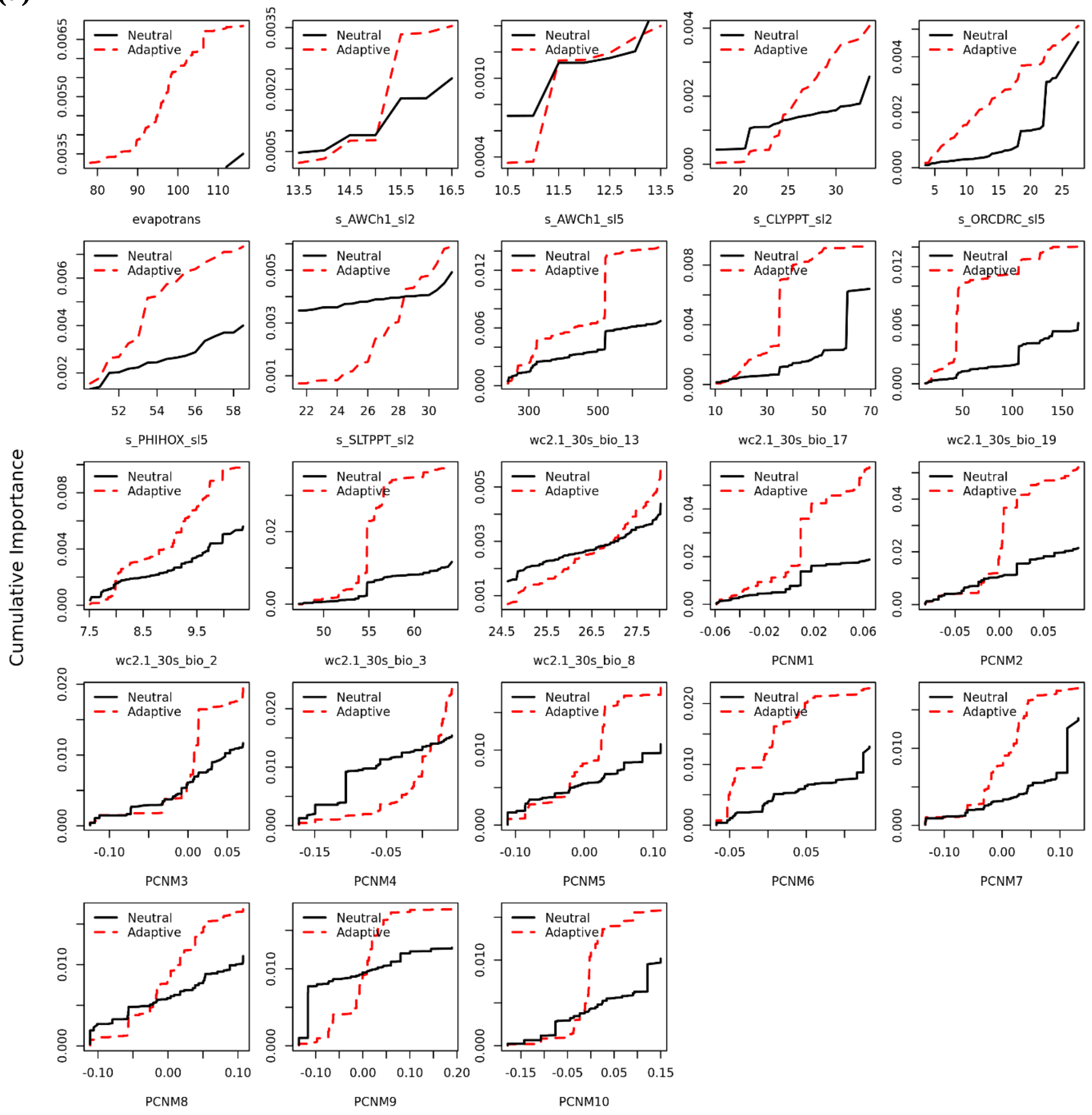

**(b)**

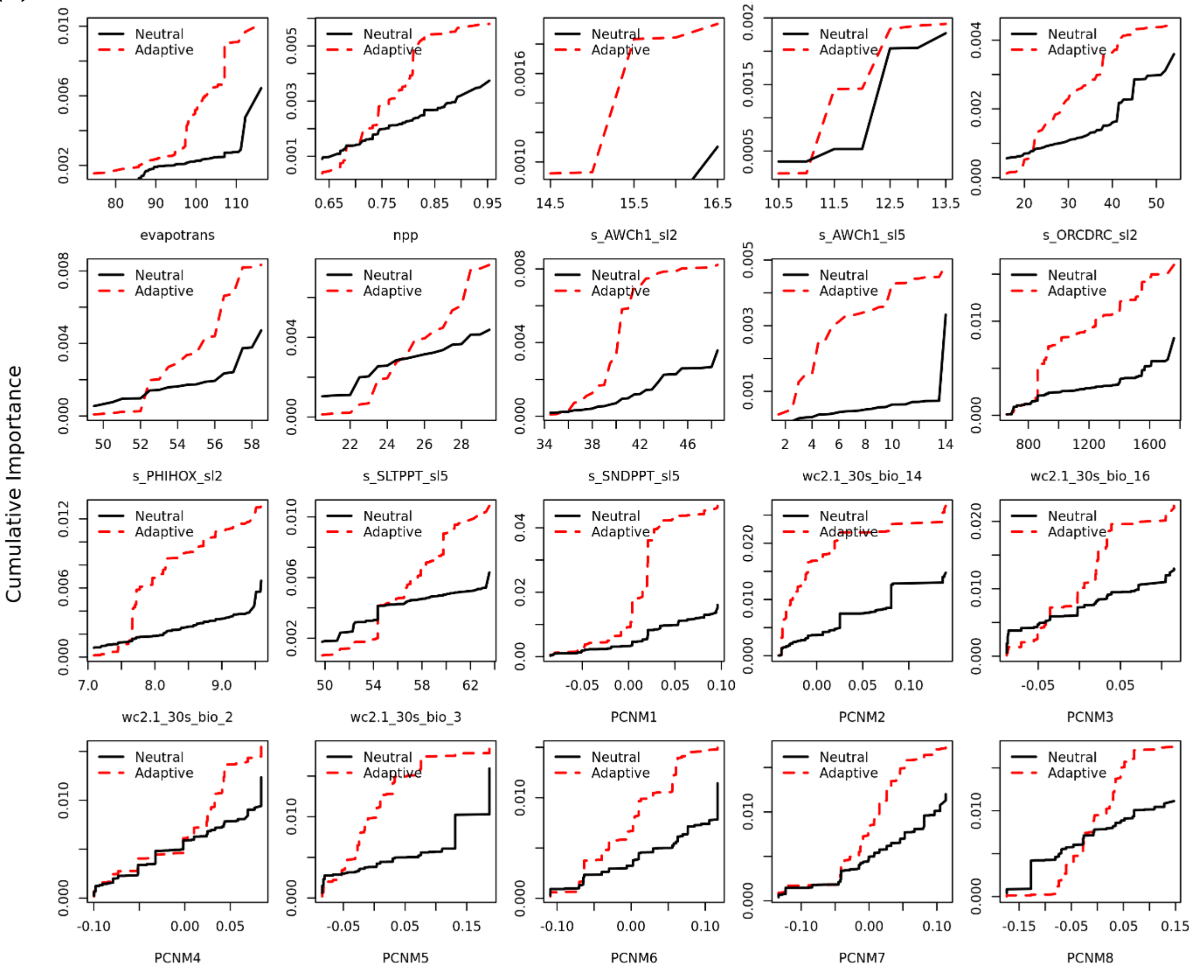
Cumulative Importance
Neutral
Adaptive
evapotrans
npp
s_AWCh1_sl2
s_AWCh1_sl5
s_ORCDRC_sl2
s_PHIHOX_sl2
s_SLTPPT_sl5
s_SNDPPT_sl5
wc2.1_30s_bio_14
wc2.1_30s_bio_16
wc2.1_30s_bio_2
wc2.1_30s_bio_3
PCNM1
PCNM2
PCNM3
PCNM4
PCNM5
PCNM6
PCNM7
PCNM8

Supplementary Figure 10.

Genetic offset of gene-environmental association (scaled between 0, lowest offset, and 1, highest offset) across the range of **(a)** *D. cochinchinensis* and **(b)** *D. oliveri* in 4 SSPs (126, 245, 370, and 585) over three bidecades (2041–2060, 2061–2080, 2081–2100) averaged across five GCMs (BCC-CSM2-MR, CNRM-ESM2-1, IPSL-CM6A-LR, MIROC6, MRI-ESM2-0.

**(a)**

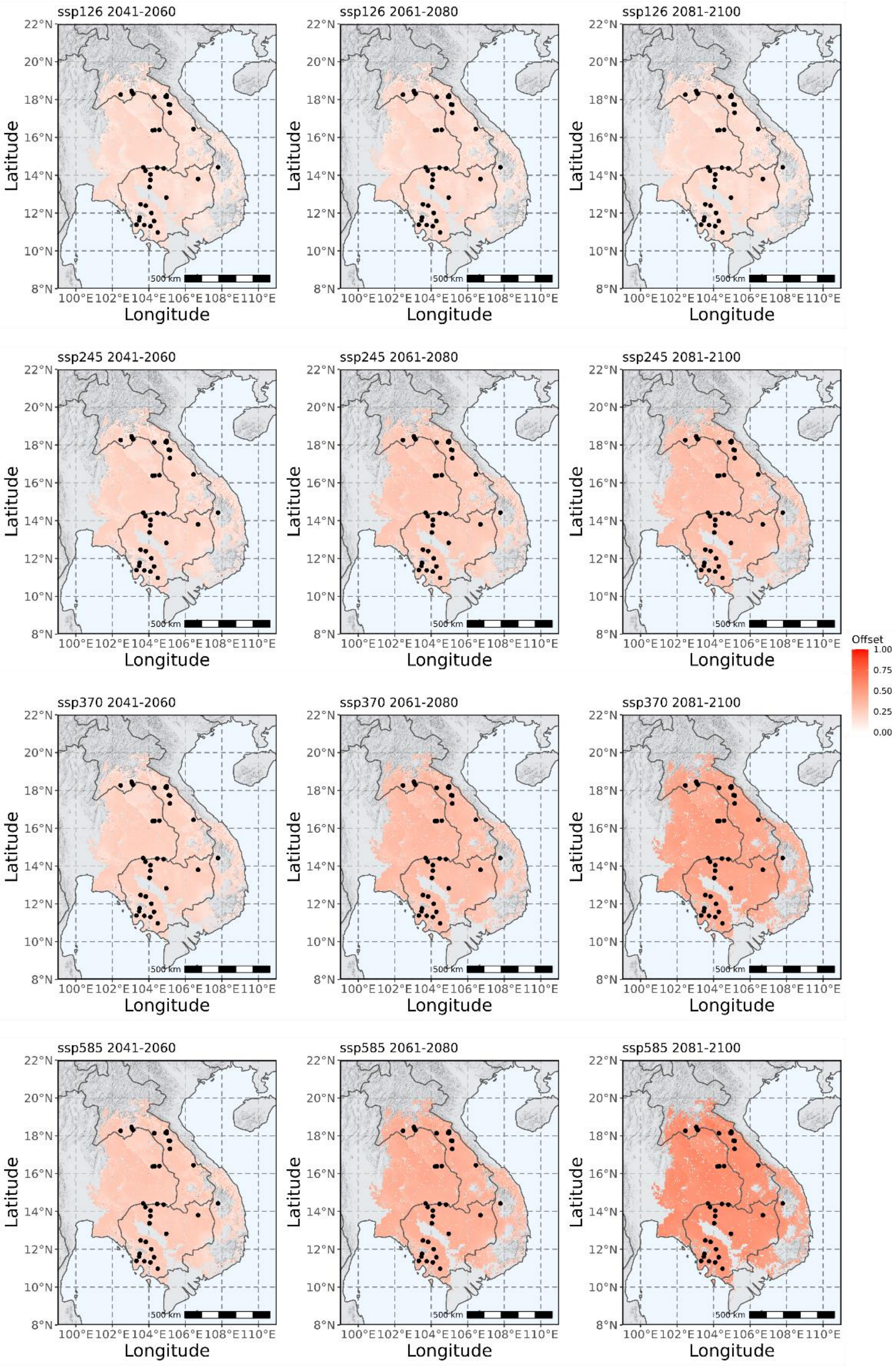

**(b)**

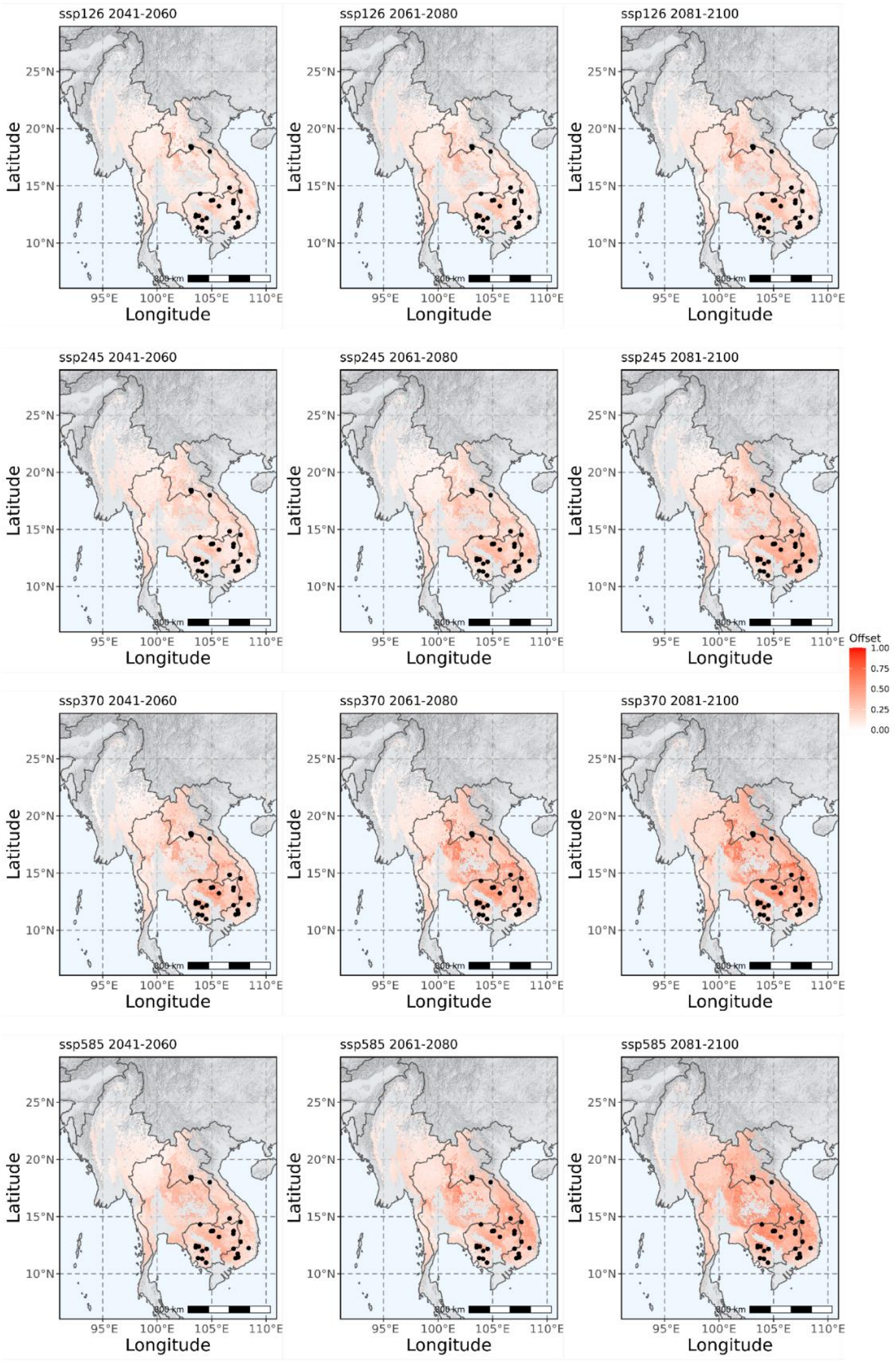
ssp126 2041-2060
ssp126 2061-2080
ssp126 2081-2100
ssp245 2041-2060
ssp245 2061-2080
ssp245 2081-2100
ssp370 2041-2060
ssp370 2061-2080
ssp370 2081-2100
ssp585 2041-2060
ssp585 2061-2080
ssp585 2081-2100
Latitude
Longitude
25°N
20°N
15°N
10°N
95°E
100°E
105°E
110°E
800 km
Offset
1.00
0.75
0.50
0.25
0.00

Supplementary Table 10.

Analysis of variance (ANOVA) table of the averaged genomic vulnerability on the interactions among time, SSP, and species.

Analysis of Variance Table

| | **Df** | **Sum Sq** | **Mean Sq** | **F-value** | **Pr (>F)** | |
|---|---|---|---|---|---|---|
| Time | 1 | 2.1558e-05 | 2.1558e-05 | 48.8719 | 2.706e-10 | *** |
| SSP | 3 | 3.9757e-05 | 1.3252e-05 | 30.0432 | 4.535e-14 | *** |
| Species | 1 | 1.0318e-04 | 1.0318e-04 | 233.9018 | < 2.2e-16 | *** |
| Time:SSP | 3 | 9.2440e-06 | 3.0810e-06 | 6.9850 | 0.0002509 | *** |
| Time:Species | 1 | 6.6060e-06 | 6.6060e-06 | 14.9762 | 0.0001902 | *** |
| SSP:Species | 3 | 1.0555e-05 | 3.5180e-06 | 7.9759 | 7.761e-05 | *** |
| Time:SSP:Species | 3 | 4.3060e-06 | 1.4350e-06 | 3.2542 | 0.0246941 | * |
| Residuals | 104 | 4.5876e-06 | 4.4100e-07 | | | |

Supplementary Table 11.

Sampling and quality statistics of genetic samples used for the ecological genomic study in *D. cochinchinensis* and *D. oliveri*.

*In a separate spreadsheet*.

Supplementary Table 12.

List of environmental data used in the landscape genomics modelling of this study.

| Symbol | Description | Category | Source | *D. cochinchinensis* | *D. oliveri* |
|---|---|---|---|---|---|
| wc2.1_30s_bio_1 | Annual mean temperature | Temperature | WorldClim | | |
| wc2.1_30s_bio_2 | Mean diurnal range | Temperature | WorldClim | V | V |
| wc2.1_30s_bio_3 | Isothermality | Temperature | WorldClim | V | V |
| wc2.1_30s_bio_4 | Temperature seasonality | Temperature | WorldClim | | |
| wc2.1_30s_bio_5 | Maximum temperature of warmest month | Temperature | WorldClim | | |
| wc2.1_30s_bio_6 | Minimum temperature of coldest month | Temperature | WorldClim | | |
| wc2.1_30s_bio_7 | Temperature annual range | Temperature | WorldClim | | |
| wc2.1_30s_bio_8 | Mean temperature of wettest quarter | Temperature | WorldClim | V | |
| wc2.1_30s_bio_9 | Mean temperature of driest quarter | Temperature | WorldClim | | |
| wc2.1_30s_bio_10 | Mean temperature of warmest quarter | Temperature | WorldClim | | |
| wc2.1_30s_bio_11 | Mean temperature of coldest quarter | Temperature | WorldClim | | |
| wc2.1_30s_bio_12 | Annual precipitation | Precipitation | WorldClim | | |
| wc2.1_30s_bio_13 | Precipitation of wettest month | Precipitation | WorldClim | V | |
| wc2.1_30s_bio_14 | Precipitation of driest month | Precipitation | WorldClim | | V |
| wc2.1_30s_bio_15 | Precipitation seasonality | Precipitation | WorldClim | | |
| wc2.1_30s_bio_16 | Precipitation of wettest quarter | Precipitation | WorldClim | | V |
| wc2.1_30s_bio_17 | Precipitation of driest quarter | Precipitation | WorldClim | V | |
| wc2.1_30s_bio_18 | Precipitation of warmest quarter | Precipitation | WorldClim | | |

| | | | | | |
|---|---|---|---|---|---|
| wc2.1_30s_bio_19 | Precipitation of coldest quarter | Precipitation | WorldClim | V | |
| wc2.1_30s_bio_elev | Digital elevation | Elevation | WorldClim | | |
| s_AwCh1_sl2 | Available soil water capacity (volumetric fraction) with FC = pF 2.0 at depth of 5 cm | Soil | SoilGrids | V | V |
| s_AwCh1_sl5 | Available soil water capacity at depth of 60 cm | Soil | SoilGrids | V | V |
| s_CLYPPT_sl2 | Clay content in percent at depth of 5 cm | Soil | SoilGrids | V | |
| s_CLYPPT_sl5 | Clay content in percent at depth of 60 cm | Soil | SoilGrids | | |
| s_ORCDRC_sl2 | Soil organic carbon stock at depth of 5 cm | Soil | SoilGrids | | V |
| s_ORCDRC_sl5 | Soil organic carbon stock at depth of 60 cm | Soil | SoilGrids | V | |
| s_PHIHOX_sl2 | Soil pH measured in water solution at depth of 5 cm | Soil | SoilGrids | | V |
| s_PHIHOX_sl5 | Soil pH measured in water solution at depth of 60 cm | Soil | SoilGrids | V | |
| s_SLTPPT_sl2 | Silt content in percent at depth of 5 cm | Soil | SoilGrids | V | |
| s_SLTPPT_sl5 | Silt content in percent at depth of 60 cm | Soil | SoilGrids | | V |
| s_SNDPPT_sl2 | Sand content in percent at depth of 5 cm | Soil | SoilGrids | | |
| s_SNDPPT_sl5 | Sand content in percent at depth of 60 cm | Soil | SoilGrids | | V |
| npp | Net primary productivity | Vegetation | The Atlas of the Biosphere | | V |

| | | | | | |
|---|---|---|---|---|---|
| evapotrans | Evapotranspiration | Vegetation | The Atlas of the Biosphere | V | V |

Supplementary Figure 11.

The user interface of the “Predict” function of *seedeR* (http://app.hung.studio/seedeR). In the left panel “Inputs”, the user can choose a species, a SSP, a time period and input the longitude and the latitude of the future restoration site. After clicking “Run”, the process will begin and print the output In the right panel “Output”.

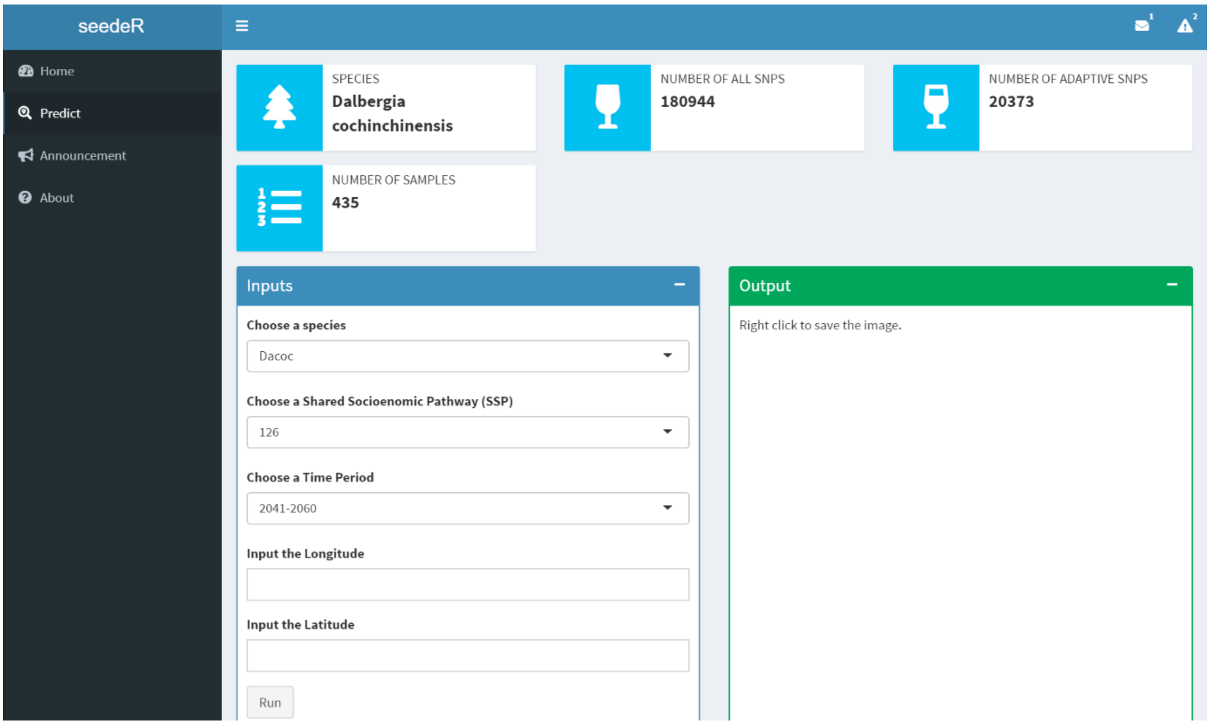
